# Amplicon structure creates collateral therapeutic vulnerability in cancer

**DOI:** 10.1101/2022.09.08.506647

**Authors:** Yi Bei, Luca Bramé, Marieluise Kirchner, Raphaela Fritsche-Guenther, Sevrine Kunz, Animesh Bhattacharya, Julia Köppke, Jutta Proba, Nadine Wittstruck, Olga A. Sidorova, Rocío Chamorro González, Heathcliff Dorado Garcia, Lotte Brückner, Robin Xu, Mădălina Giurgiu, Elias Rodriguez-Fos, Richard Koche, Clemens Schmitt, Johannes H. Schulte, Angelika Eggert, Kerstin Haase, Jennifer Kirwan, Anja I.H. Hagemann, Philipp Mertins, Jan R. Dörr, Anton G. Henssen

**Author notes:** These authors jointly supervised this work (Jan R. Dörr, Anton G. Henssen). Correspondence should be addressed to A.G.H. and J.R.D.

## Abstract

Although DNA amplifications in cancers frequently harbor passenger genes alongside oncogenes, the functional consequence of such co-amplifications and their impact for therapy remains ill-defined. We discovered that passenger co-amplifications can create amplicon structure-specific collateral vulnerabilities. We present the DEAD-box helicase 1 (*DDX1*) gene as a *bona fide* passenger co-amplified with *MYCN* in cancers. Survival of cancer cells with *DDX1* co-amplifications strongly depends on the mammalian target of rapamycin complex 1 (mTORC1). Mechanistically, aberrant DDX1 expression inhibits the tricarboxylic acid cycle through a previously unrecognized interaction with dihydrolipoamide S-succinyltransferase, a component of the alpha-ketoglutarate dehydrogenase complex. Cells expressing aberrant DDX1 levels compensate for the metabolic shift by enhancing mTORC1 activity. Consequently, pharmacological mTORC1 inhibition triggered cell death specifically in cells harboring the *DDX1* co-amplification. This work highlights a significant contribution of passenger gene alterations to the therapeutic susceptibility of cancers.

**Graphical abstract:** 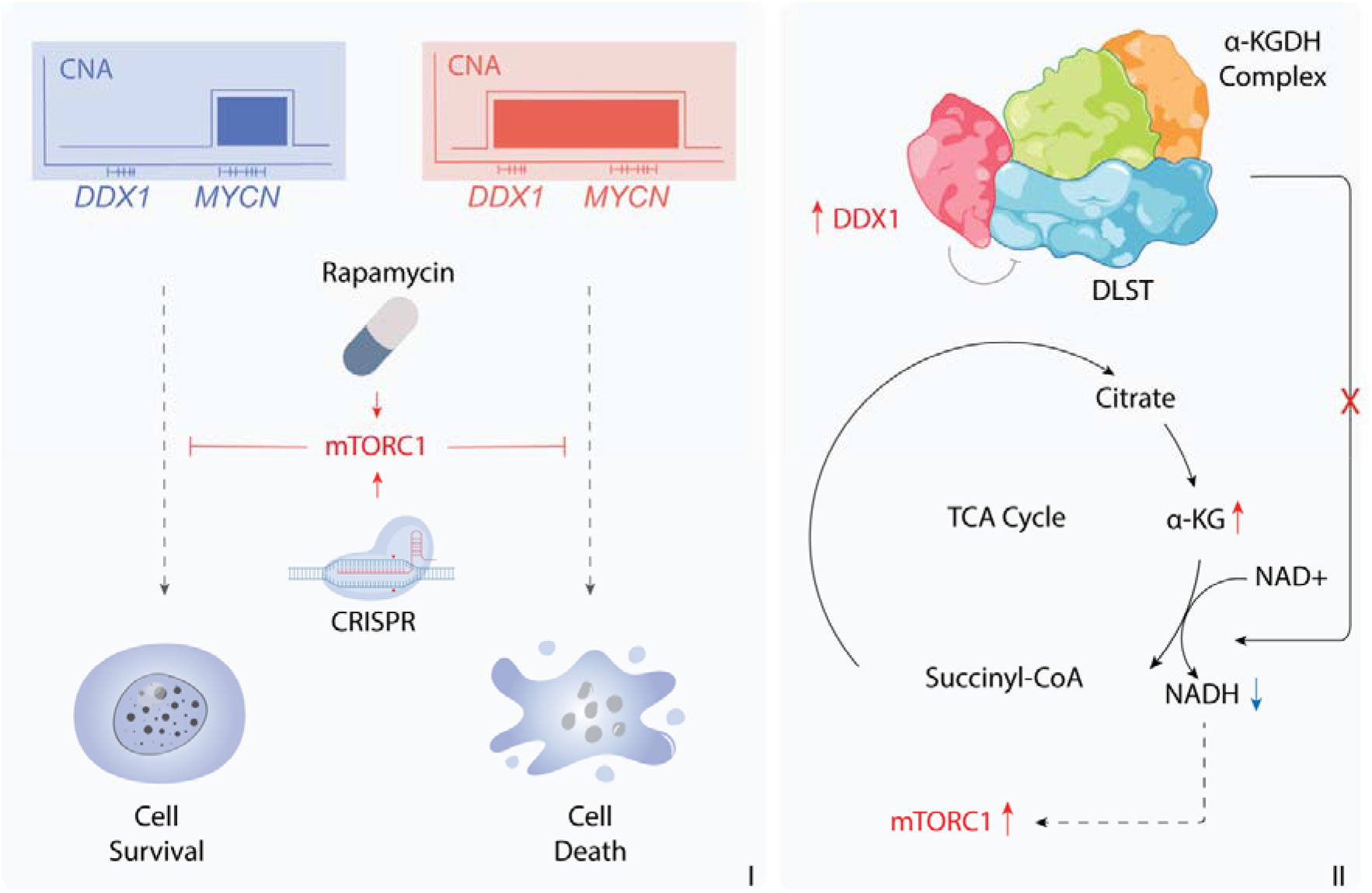

## Introduction

Somatic DNA amplification is a common phenomenon in cancers and one of the most important causes of excessive oncogene expression (Beroukhim et al., 2010; Calabrese et al., 2020; Schwab, 1999). Recent cancer genome sequencing efforts have revealed fundamental insights into amplicon structures (Helmsauer et al., 2020; Kim et al., 2020; Morton et al., 2019; Rosswog et al., 2021; Shoshani et al., 2021). Accordingly, DNA amplification exists in at least two forms: (i) self-repeating arrays on a chromosome (homogeneously staining regions, HSR) and (ii) many individual circular extrachromosomal DNA molecules (extrachromosomal DNA, ecDNA)(Shimizu et al., 1998; Turner et al., 2017; van Leen et al., 2022; Verhaak et al., 2019; Yi et al., 2022). The genomic boundaries of such amplicons are not randomly distributed around oncogenes but are largely defined by the location of nearby core-regulatory enhancer elements (Helmsauer *et al*., 2020; Morton *et al*., 2019). Enhancers that are included on amplicons are required for sustained oncogene expression (Helmsauer *et al*., 2020; Morton *et al*., 2019; Wu et al., 2019; Zhu et al., 2021), suggesting that genomic regions co-amplified with oncogenes are under strong positive selection. One prominent example of extrachromosomal oncogene amplification is the *MYCN-*harboring amplicon in neuroblastoma, which can carry multiple enhancers and can include many megabases of additional genomic sequences (Helmsauer *et al*., 2020). These emerging structural properties of DNA amplicons may explain the recurrently observed co-amplification of passenger genes in the vicinity of the amplified oncogene (Albertson, 2006; Chen et al., 2014; Schwab, 1998; Scott et al., 2003). The fact that such passenger genes are retained on amplicons in cancer cells implies that their amplification does not compromise cancer cell fitness. Whether and how passenger gene co-amplifications alter cancer cell physiology and therapeutic susceptibility, however, has not been investigated conclusively to date.

Inspired by the concept of collateral lethality, which has been employed to identify cancer-specific therapeutic vulnerabilities resulting from co-deletions of genes neighboring tumor suppressor genes (Dey et al., 2017; Muller et al., 2015; Muller et al., 2012), we here investigated whether passenger co-amplification could present novel amplicon structure-specific therapeutic strategies for tumors harboring oncogene amplifications, for which drug development has remained largely elusive to date.

## Results

### Passenger genes are frequently co-amplified with oncogenes in cancer

The recent discovery that oncogene amplifications encompass large neighboring genomic regions with regulatory elements suggests that co-amplification of nearby passenger genes may be more common in cancers than previously anticipated. To assess the frequency of passenger gene co-amplification, we analyzed whole-genome sequences from well-characterized cohorts of childhood and adult tumors from the Pan-Cancer Analysis of Whole Genomes (PCAWG) study (Pan-cancer analysis of whole genomes, 2020) and Tumor Alteration Relevant for Genomics-driven Therapy (TARGET) database (Pugh et al., 2013; Van Allen et al., 2014). We classified amplified genes as oncogenes or passenger genes based on the Catalogue Of Somatic Mutations In Cancer (COSMIC) cancer annotation (Sondka et al., 2018). Passenger gene amplification was common in all analyzed tumor entities (Figure 1A and Figure S1A-S1B), but very rarely occurred on amplicons not harboring oncogenes. Intriguingly, almost all oncogene-containing amplicons also harbored passenger genes (Figure S1C-S1E). Thus, passenger gene co-amplification is common in cancers, which may affect the physiology of cancer cells and generate collateral therapeutic vulnerabilities.

**Figure 1.**
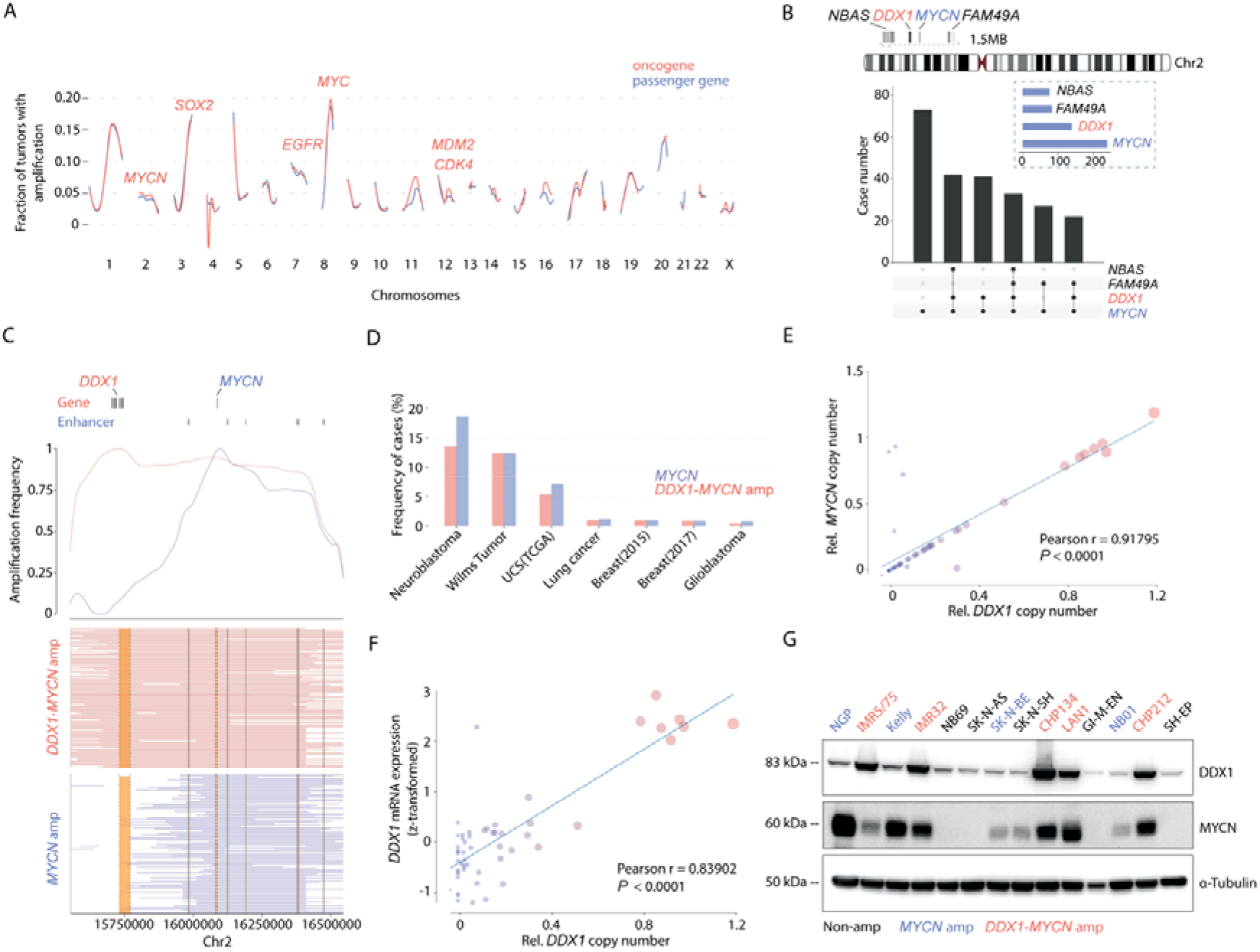
Passenger genes are frequently co-amplified with oncogenes across tumor entities. **A**, Fraction of tumors (*N* = 2960) from PCAWG and TARGET datasets with oncogene amplifications (red) or passenger gene amplifications (blue) throughout the genome fitted by local regression (LOESS). The annotated genes are amongst the most commonly altered oncogenes in cancers. **B**, Chromosome 2 schematic highlighting the area of *MYCN* amplification and passenger genes recurrently included on the amplicon (top). Upset plot (bottom) for the co-amplification frequency of three passenger genes, *NBAS, DDX1* and *FAM49A*, identified on the *MCYN* amplicon in a cohort of 556 neuroblastomas. **C**, Density plot for the amplification frequency near *MYCN*, measured using copy number profiles from *238 MYCN*-amplified neuroblastoma patients with (red) and without (blue) *DDX1* co-amplification. **D**, Frequency of *MYCN* amplifications with (red) and without (blue) *DDX1* co-amplification in different tumor entities. UCS, uterine carcinosarcoma; Lung cancer, lung adenocarcinoma. Breast 2015, BRCA_igr_2015; Breast 2017, BRCA_mbcproject_wagle_2017. **E**, Correlation between *DDX1* copy number and *MYCN* copy number derived from the TARGET neuroblastoma dataset (Pearson r = 0.91795, *P* < 0.0001, *N* = 59). Size of dots reflects the relative *MYCN* and *DDX1* copy number. **F**, Correlation between *DDX1* copy number and DDX1 mRNA expression derived from the TARGET neuroblastoma dataset (microarray) (Pearson r = 0.83952, *P* < 0.0001, *N* = 59). **G**, Western immunoblot of DDX1 and MYCN in a panel of neuroblastoma cell lines with (red) and without (blue) *DDX1-MYCN* co-amplifications, compared to cell lines without *MYCN* amplifications (black).

### DDX1 is frequently co-amplified with MYCN in cancers

To uncover collateral vulnerabilities induced by passenger gene co-amplification, we focused our analysis on *MYCN* amplifications, which are frequent in many tumor entities, particularly neuroblastoma, and are often associated with high-risk disease and poor therapeutic outcome (Brodeur et al., 1984; Helmsauer *et al*., 2020; Koche et al., 2020; Maris, 2010; Seeger et al., 1985; Weiss et al., 1997). As a basic helix-loop-helix oncogenic transcription factor, *MYCN* remains inapproachable for direct therapeutic interventions (Chen et al., 2018), making it an ideal and clinically highly relevant candidate to test our hypothesis. Analysis of 556 published neuroblastoma genome-wide copy number profiles (Depuydt et al., 2018a; Depuydt et al., 2018b) identified 238 neuroblastomas with *MYCN* amplifications (Figure 1B). In line with our previous reports (Helmsauer *et al*., 2020), the *MYCN* amplicon on average encompassed a large 1-3 Mb region with several co-amplified passenger genes, including *DDX1, NBAS*, and *FAM49A. DDX1*, a gene encoding for the Asp-Glu-Ala-Asp (DEAD) functioning DNA:RNA ATPase DDX1 (Godbout et al., 1998; Linder et al., 1989; Schmid and Linder, 1992; Wassarman and Steitz, 1991), was the most recurrently co-amplified passenger gene with *MYCN* in neuroblastomas (57.98%, 138 out of 238) (Figure 1B, 1C and S1E). No *DDX1* amplifications without *MYCN* were detectable in 556 cancer genomes. *DDX1-MYCN* co-amplification also occurred in several other cancer entities (Figure 1D). Combined analysis of copy number and mRNA expression of *DDX1* and *MYCN* confirmed a significant positive correlation in expression and aberrantly high *DDX1* expression levels in the context of co-amplification (Figure 1E and Figure S2). Consistently, neuroblastoma cell lines harboring a *DDX1*-*MYCN* co-amplification had elevated DDX1 protein and mRNA levels compared to those lacking *DDX1* co-amplifications or cells without *MYCN* amplification (Figure 1F, 1G and Figure S2). Thus, *DDX1*-*MYCN* co-amplification is present in a considerable fraction of cancers and is associated with aberrantly high DDX1 expression, which could affect cancer cell physiology.

### DDX1 is a bona fide passenger gene

In the classic dichotomous model of driver and passenger genes, passenger genes are defined as genetic moieties that are altered in their expression or sequence but unlike oncogenes do not drive cancer initiation or progression (Greenman et al., 2007). Although DDX1 has been implicated in many critical cellular activities, such as RNA regulation (Chen et al., 2002; Han et al., 2014) and DNA damage repair (Li et al., 2008), evidence for its oncogenic potential is scarce. To investigate the role of DDX1 in the pathogenesis of neuroblastoma in an *in vivo* experimental system, we generated a transgenic zebrafish line that stably expresses human DDX1 in the peripheral sympathetic nervous system under the control of the zebrafish dopamine-β-hydroxylase gene (*dβh*) promoter. Transgenic fish were created by injection of *dβh-DDX1:CryAA-mCerulean* into zebrafish fertilized eggs (Figure 2A). Transgenic integration was identified in fish (generation F0) by fluorescent reporter expression and human DDX1 expression was confirmed in offspring (F1) by immunoblotting (Figure 2B and 2C). No tumors developed with transgenic expression of DDX1 alone in fish of F1 generation with stable transgenic germline integration (Figure 2B, 2C and 2D), indicating that DDX1 cannot drive tumorigenesis. Given the strong association of DDX1 and MYCN expression due to co-amplification in human neuroblastoma (Figure 1), we next tested whether high levels of DDX1 expression could cooperate with MYCN to affect the onset or penetrance of neuroblastic zebrafish tumors. An established zebrafish model that expresses human MYCN under control of the *dβh* promoter was interbred with DDX1-expressing zebrafish (Tao et al., 2017). Neuroblastic tumors in the adrenal gland analogue developed in 100% of the *dβh-MYCN*; *dβh-DDX1* progeny by 8 weeks of age, compared to an overall penetrance of 97% for the fish with MYCN expression alone, which also developed around the same time (Figure 2D and Figure S3A). Thus, high levels of DDX1 expression does not significantly alter the initiation or progression of MYCN-driven neuroblastic tumors *in vivo*.

**Figure 2.**
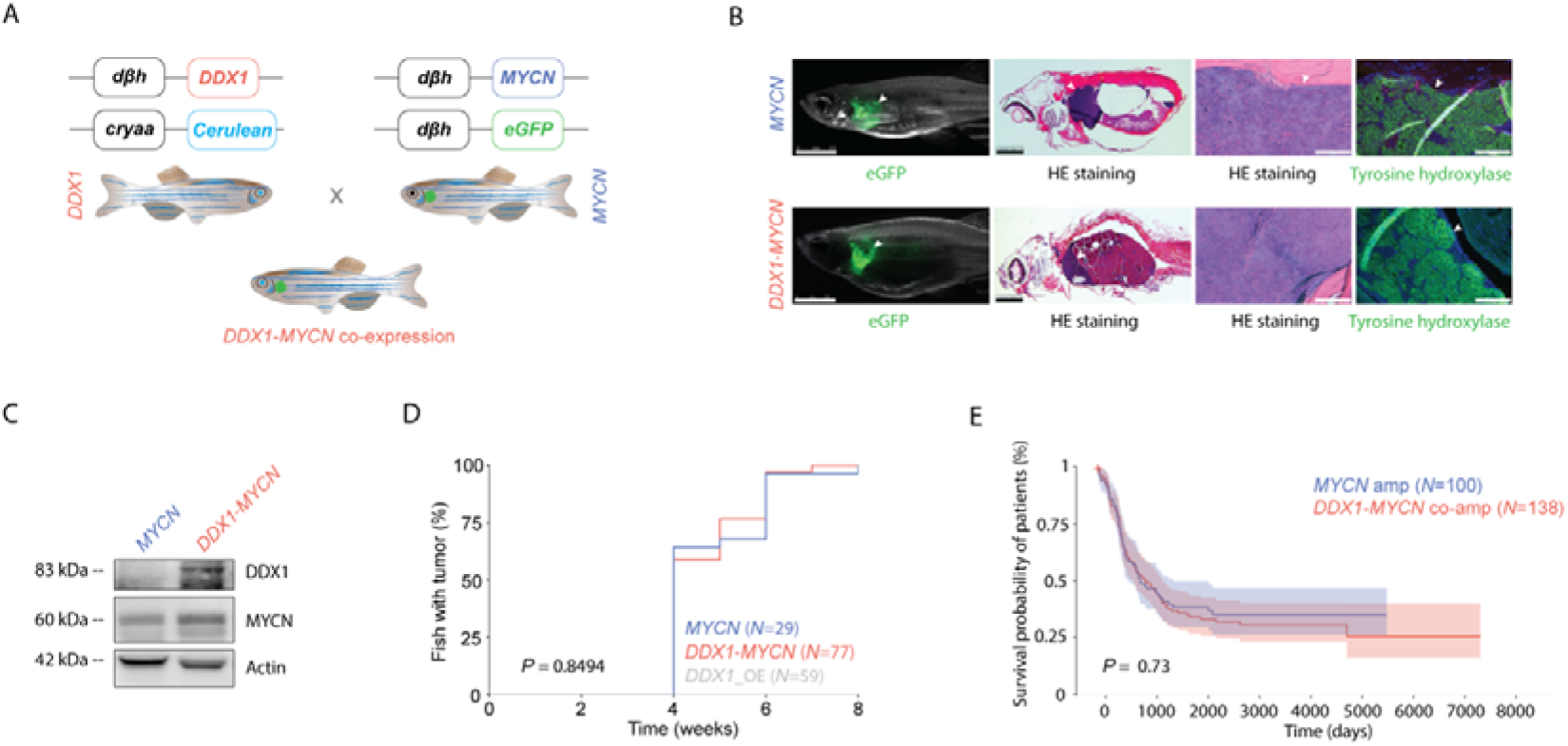
Ectopic DDX1 expression does not alter MYCN-driven tumorigenesis in zebrafish. **A**, Schematic figure showing the generation of DDX1-MYCN co-expressing zebrafish through breeding of tg (*dβh*-*DDX1*:*CryAA*-*mCerulean*) and tg (*dβh*-*MYCN*: *dβh*-*eGFP*) zebrafish. **B** From left to right, exemplary images from transgenic zebrafish tg (*dβh*-*DDX1*:*CryAA*-*mCerulean*) and tg (*dβh*-*MYCN*: *dβh*-*eGFP*) green fluorescent neuroblastic tumors in the adrenal medulla analogue (interrenal gland, white arrowhead). Hematoxylin & Eosin (HE) staining of sagittal paraffin sections from the same fish. Magnification into the tumor area from sections shown left, with HE and tyrosine hydroxylase (green) staining. Scale bar from left to right: 2.5 mm, 1 mm, 100 µm and 100 µm. Arrowheads point to neuroblastic tumors in zebrafish. **C**, Western immunoblot of DDX1 and MYCN in zebrafish tumor cell extracts. **D**, Cumulative frequency of neuroblastic tumors in stable transgenic zebrafish by Kaplan-Meier analysis (*DDX1-MYCN* vs. *MYCN*, Kolmogorow-Smirnow-Test, *P* = 0.8494). **E**, Kaplan-Meier analysis of patients with *DDX1*-*MYCN* co-amplification compared to patients with *MYCN* amplifications lacking *DDX1* co-amplification (Log-Rank Test, *P* = 0.73).

In line with our observation in zebrafish, *DDX1-MYCN* co-amplification in human neuroblastoma was not associated with differences in overall patient survival compared to patients with tumors only harboring *MYCN* amplifications (Figure 2E), indicating that DDX1 does not significantly alter clinically relevant malignant features of neuroblastoma. To further test the effect of high DDX1 expression in cancer cells, we selected human neuroblastoma cell lines harboring *MYCN* amplifications, not including *DDX1* and introduced a doxycycline-inducible DDX1 expression vector. Ectopic expression of DDX1 did not affect neuroblastoma cell proliferation (Figure S3B-S3E). In line with its role as a passenger gene, short hairpin RNA (shRNA) mediated DDX1 knockdown in *DDX1*-*MYCN* co-amplified neuroblastoma cell lines did not reduce neuroblastoma proliferation (Figure S3F-S3I). Although DDX1 did not influence the tumorigenic properties of neuroblastoma cells, ectopic DDX1 expression was associated with significantly reduced neuroblastoma cell size (Figure S3J-S3K). This suggests that DDX1 acts as a *bona fide* passenger gene in neuroblastoma, but that its aberrant expression influences cellular physiology. This raises the possibility that the non-tumorigenic effects of *DDX1*-*MYCN* co-amplification could generate collateral lethal dependencies.

### DDX1 co-amplification is accompanied by a collateral mTORC1 dependency

Having confirmed that *DDX1* acts as a *bona fide* passenger gene in *MYCN*-amplified neuroblastoma and that its amplification is structurally linked to *MYCN*, we asked whether its aberrantly high expression could result in collateral genetic dependencies. To identify such dependencies, we analyzed copy number profiles of human cancer cell lines from the Broad-Novartis Cancer Cell line Encyclopedia (CCLE)(Barretina et al., 2012; Ghandi et al., 2019) and selected all cancer cell lines with *MYCN* amplification. Next, we compared the genetic dependencies of cell lines with *DDX1-MYCN* co-amplification to those only harboring *MYCN* amplifications by analyzing genome-scale pooled CRISPR/Cas9 loss of viability screens in over 700 genomically characterized human cancer cells from 26 tumor lineages as part of the Cancer Dependency Map (Dempster et al., 2019; Meyers et al., 2017). We confirmed that *MYCN* copy number and expression levels were comparable between these two groups (Figure S4A and S4B). We calculated the dependency score as well as its median difference for each gene between cell lines with *MYCN* amplifications vs. cell lines with *DDX1-MYCN* co-amplifications (Figure 3A). This revealed that *DDX1*-*MYCN* co-amplification was significantly associated with high genetic dependency to mTORC1 complex members mTOR and its scaffold protein RAPTOR, which plays an important role in mTORC1 activation (Carrière et al., 2008; Yao et al., 2016) (Figure 3B, Figure S4C and Table S1). In line with an increased RAPTOR dependency in the context of *DDX1*-*MYCN* co-amplification, dependency scores for RAPTOR significantly negatively correlated with the *DDX1* copy number in *MYCN*-amplified neuroblastoma cell lines (Figure 3C, Pearson coefficient = −0.5996, *P* = 0.0152, Figure S4D and S4E). Moreover, ectopic expression of DDX1 in *MYCN*-amplified cell lines was sufficient to increase RAPTOR dependency, as evidenced by reduced clonogenicity of cells after CRISPR-Cas9-mediated knockout of RAPTOR (Figure 3D, 3E and Figure S4F and S4G). This indicates that high DDX1 expression, as observed in the context of *DDX1*-*MYCN* co-amplification, can generate an mTORC1 dependency.

**Figure 3.**
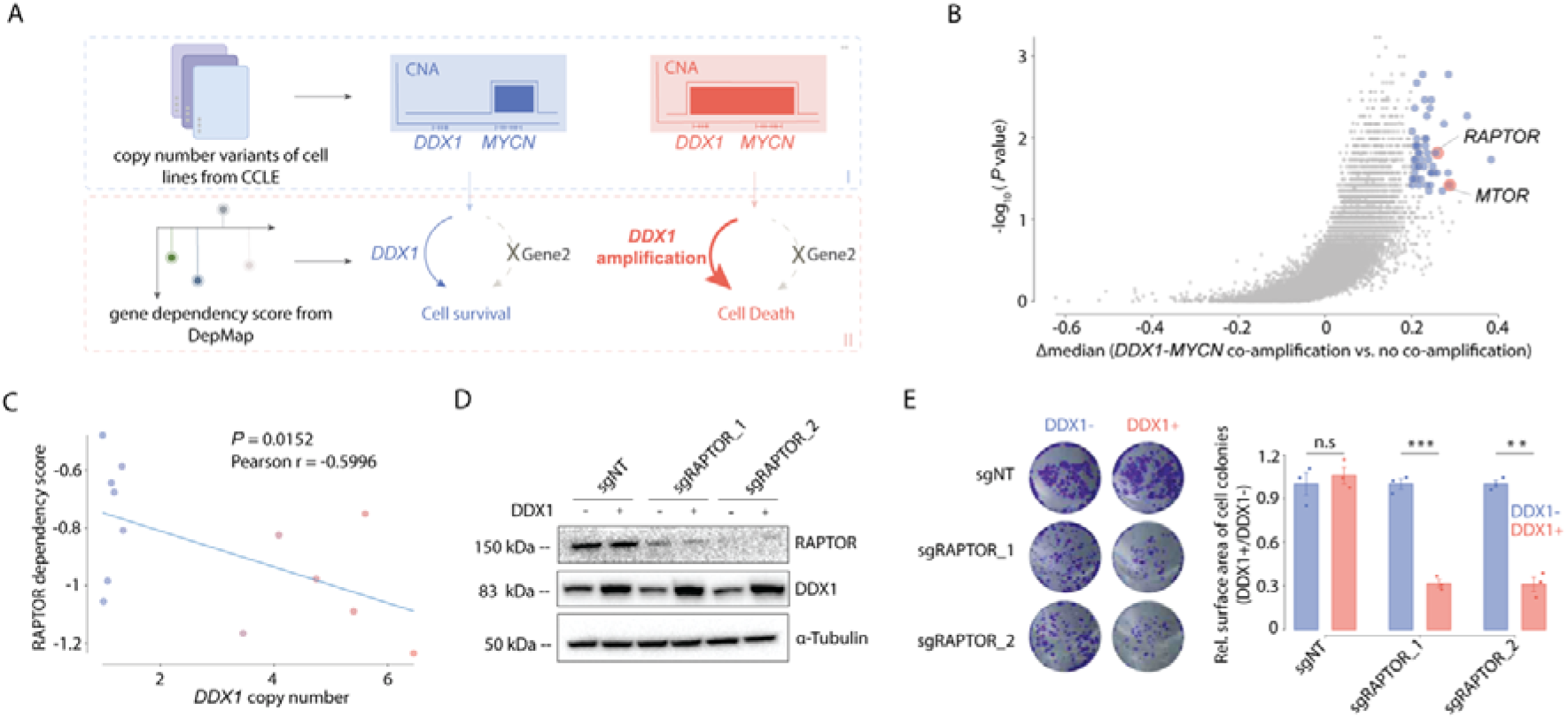
Neuroblastoma cells harboring *DDX1-MYCN* amplifications depend on mTORC1. **A**, Schematic figure illustrating the approach for the identification of collateral lethal dependencies in *DDX1*-*MYCN* co-amplified cancer cells. *DDX1* and *MYCN* copy numbers across cancer cell lines were retrieved from the Cancer Cell Line Encyclopedia (CCLE II). CRISPR-Cas9-based gene dependency scores from DepMap of cell lines with *MYCN*-amplifications were compared to those with *DDX1-MYCN* co-amplification. **B**, Difference in gene dependency scores between cancer cell lines with *DDX1*-*MYCN* co-amplification vs. cell lines with *MYCN* amplifications compared to the log-transformed *P* values (Wilcoxon; candidate collateral lethal dependencies in *DDX1*-*MYCN* co-amplified cancer cells, blue; mTORC1 complex, blue). **C**, Correlation between *DDX1* copy number and dependency scores (CERES) for *RAPTOR* in neuroblastoma cell lines (Pearson correlation analysis, *R* = −0.5996, *P* = 0.0152, *N* = 13). **D**, Western immunoblot of RAPTOR and DDX1 in the KELLY cells transduced with the doxycycline-inducible DDX1-mcherry vectors and with two pairs of sgRNAs targeting *RAPTOR (*sgRAPTOR) or a non-targeting sgRNA (sgNT) as well as Cas9 in the presence and absence of doxycycline (1 µg/ml). Tubulin serves as a loading control. **E**, Representative images of cell colonies formed by KELLY cells transduced with the doxycycline-inducible DDX1-mcherry vectors and with two pairs of sgRNA targeting *RAPTOR (*sgRAPTOR) or non-target sgRNA (sgNT) as well as Cas9 in the presence and absence of doxycycline (1 µg/ml) and stained with crystal violet (left). Quantification of colony numbers (right, mean ± s.e. *N* = 3 biological replicates; Welch t-test, *P* = 0.564, 0.000117 and 0.00131 for sgNT, sgRAPTOR_1 and sgRAPTOR_2, respectively).

### DDX1 overexpression is sufficient to induce mTOCR1 pathway activation

To understand the mechanism of DDX1-induced mTORC1 dependency, we analyzed previously published neuroblastoma gene expression data from 709 patients (Oberthuer et al., 2015). Intriguingly, high DDX1 expression was significantly associated with gene expression programs linked to high mTORC1 pathway activation in primary neuroblastomas (Figure 4A and 4B, q = 0.01, NES = 1.675, Table S2). To test whether DDX1 expression was sufficient to induce mTORC1 pathway activation, we analyzed mTORC1 activity by RNA sequencing and immunoblot analyses of *MYCN*-amplified neuroblastoma cells after ectopic DDX1 expression. Indeed, ectopic DDX1 expression was accompanied by significant differential expression of genes associated with mTORC1 pathway activation (Figure 4C and 4D, q = 0.028, NES = 2.09, Table S3 and Figure S5A). Furthermore, phosphorylation of mTOR at Ser2448 and P70-S6K at Thr389, signs of mTORC1 pathway activation (Chiang and Abraham, 2005; Hoeffer and Klann, 2010; Takei and Nawa, 2014; Xiao et al., 2009), increased in neuroblastoma cells after ectopic DDX1 expression (Figure 4E, Figure S5B and S5C). In turn, shRNA-mediated DDX1 knockdown in *DDX1*-*MYCN* co-amplified neuroblastoma cells resulted in reduced phosphorylation of mTOR and P70-S6K (Figure 4F and Figure S5D and S5E). This suggests that DDX1 is sufficient to drive mTORC1 pathway activation in the context of *MYCN* amplification and could thereby generate a dependency on mTORC1 in cancer cells harboring *DDX1-MYCN* amplifications.

**Figure 4.**
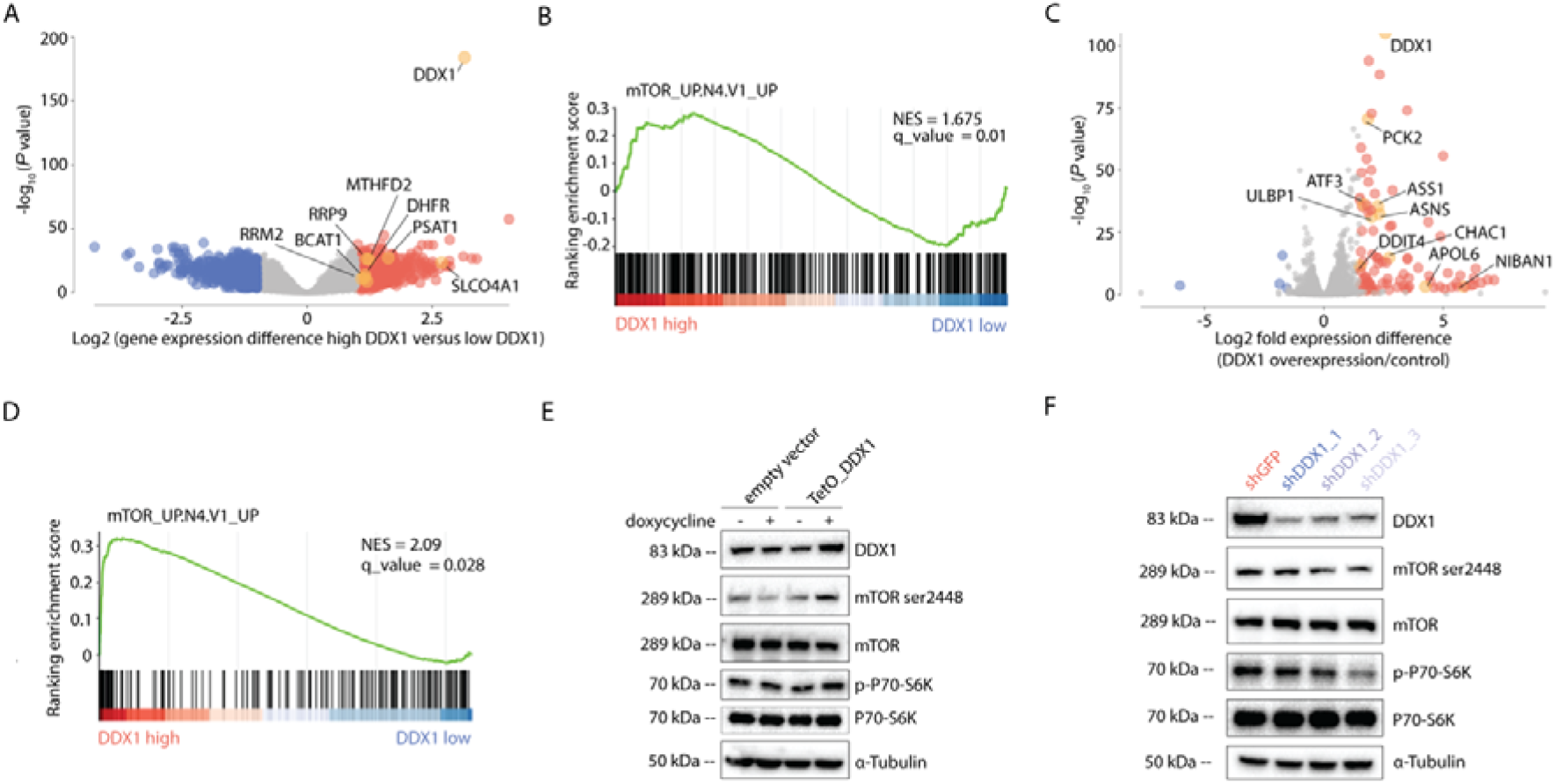
High DDX1 expression results in mTOCR1 pathway activation. **A**, Volcano plot of genes differentially expressed between primary neuroblastomas with high vs. low *DDX1* mRNA expression (*N* = 709 patients; genes with significantly lower expression, blue; genes with significantly higher expression, red, genes known to be regulated by mTORC1 signaling, orange). **B**, Gene set enrichment analysis (GSEA) based on a set of genes regulated by mTORC1 measured in genes differentially expressed in tumors with high vs. low DDX1 expression. **C**, Volcano plot of genes differentially expressed in KELLY cells with vs. without ectopic DDX1 expression (*N* = 3 independent replicates; genes with significantly lower expression, blue; genes with significantly higher expression, red, genes known to be regulated by mTORC1 signaling, orange). **D**, GSEA based on a set of genes regulated by mTORC1 measured in genes differentially expressed in KELLY cells harboring a *MYCN* amplification with vs. without ectopic DDX1 expression. **E**, Western immunoblot of mTOR ser2448 phosphorylation and P70-S6K Thr389 phosphorylation in KELLY cells 48h after inducible expression of DDX1. KELLY cell transduced with empty vector serve as negative controls. **F**, Western immunoblot of mTOR ser2448 phosphorylation and P70-S6K Thr389 phosphorylation in IMR5/75 cell after shRNA-mediated knock down of DDX1 using three independent shRNAs (blue) compared to cells expressing shRNA targeting GFP (red).

### DDX1 interacts with alpha-KGDH complex members

To investigate the mechanism by which DDX1 induces mTORC1 pathway activation, we performed immunoprecipitation of DDX1 followed by mass spectrometry-based proteomics in two *MYCN*-amplified neuroblastoma cell lines with and without *DDX1* co-amplification to identify proteins that associate with DDX1 in the context of high DDX1 expression, respectively (Figure 5A, 5B). In addition to known interactors of DDX1, e.g., eIF4G2, FAM98B and c14orf166 (Pérez-González et al., 2014), three members of the α-KGDH complex DLST, OGDH, and DLD were significantly enriched after DDX1 immunoprecipitation, particularly in cells with *DDX1-MYCN* co-amplification (Figure 5C, and Table S4). The interaction of these proteins was confirmed by co-immunoprecipitation followed by immunoblotting (Figure 5D). The α-KGDH complex predominantly localizes to the mitochondria and critically regulates electron transport chain activity and tricarboxylic acid cycle (TCA) flux. Even though DDX1 is mostly localized in the cytoplasm and nucleus in normal cells, it can associate with mitochondria during embryonal development and immune activation (Wang et al., 2022; Zhang et al., 2011). To investigate the localization of DDX1 in neuroblastoma cells, we generated cell lines expressing DDX1 fused to mCherry. Indeed, ectopically expressed DDX1-mCherry significantly co-localized with thiol-reactive chloromethyl fluorescently labeled mitochondria (Figure 5E). Thus, DDX1 can interact with α-KGDH complex members, especially when expressed at supraphysiologic levels.

**Figure 5.**
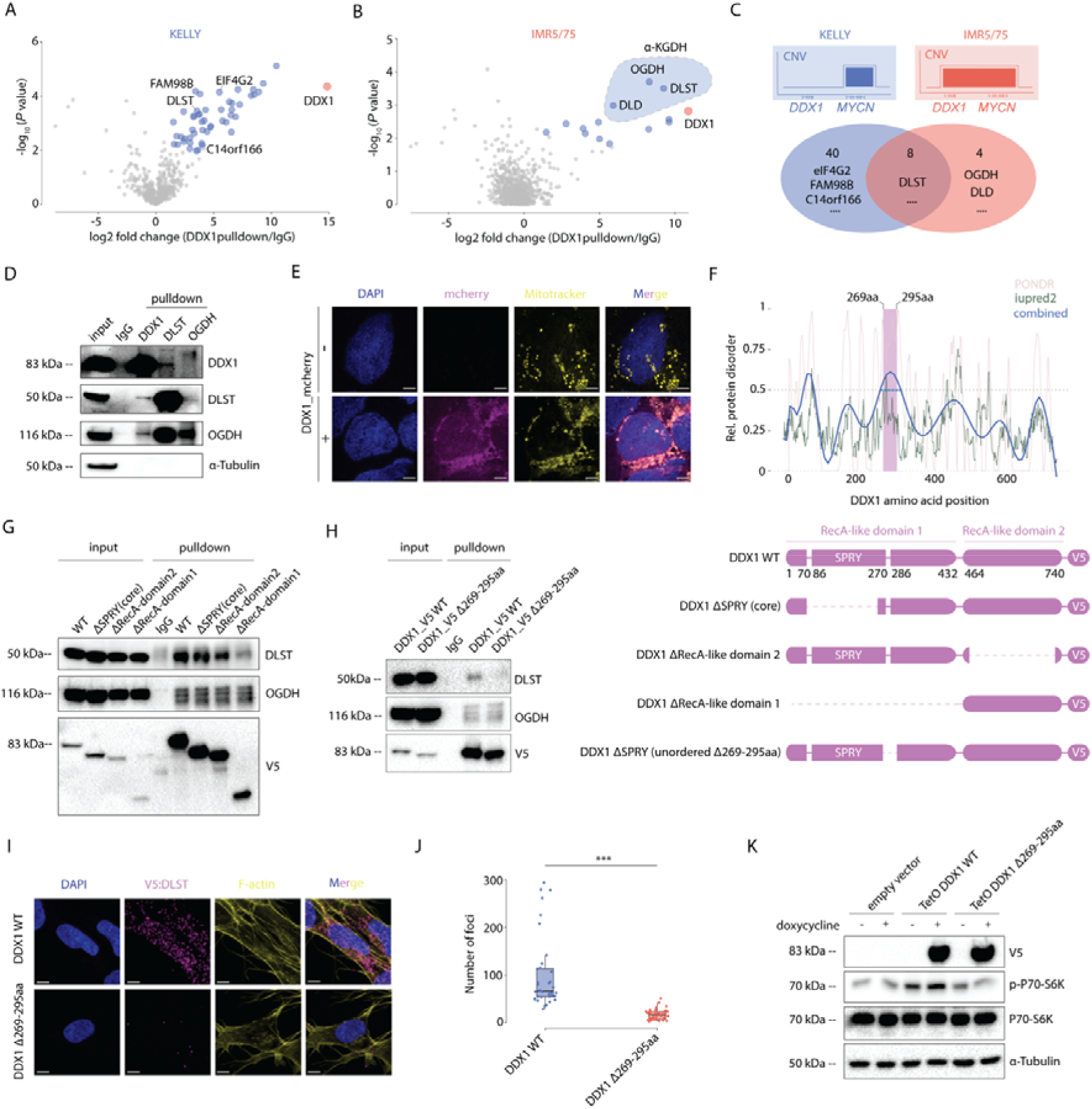
DDX1 interacts with α-KGDH complex members and its interaction is required for mTORC1 pathway activation. **A-B**, Volcano plot of proteins significantly enriched after immunoprecipitation of DDX1 in KELLY cells harboring *MYCN* amplifications without *DDX1* co-amplifications (**A**) and in IMR5/75 cells harboring *DDX1-MYCN* co-amplifications (**B**) measured using LC-MS/MS (significantly enriched proteins, blue; DDX1 marked in red; dotted line with blue filling marks α-KGDH complex members). **C**, Schematic of the amplicon structure in KELLY and IMR5/75 cells (top). Venn diagram (bottom) of the proteins identified after immunoprecipitation of DDX1 in KELLY cells lacking *DDX1* co-amplifications compared to IMR5/75 cells harboring *DDX1-MYCN* co-amplifications. **D**, Western immunoblot of DDX1, DLST, OGDH and α-tubulin in IMR5/75 protein extracts before and after immunoprecipitation using antibodies directed against DDX1, DLST, OGDH or non-specific immunoglobulins (IgG). **E**, Representative confocal fluorescence imaging photomicrographs of KELLY cells expressing DDX1-mCherry (magenta), in which mitochondria were stained by MitoTracker DeepRed (yellow) and the nucleus is stained by DAPI (blue; scale bar: 6 µm). **F**, Prediction of disordered regions in DDX1 (top) using Predictor of Natural Disordered regions (PONDR, XL1_XT, pink), Iupred2(green) and polynomial fitted model (blue; the position of amino acids 269-295 is marked in purple). Schematic illustration (bottom) of protein domains in DDX1 as well as engineered DDX1 mutants (DDX1-ΔSPRY (core), Δ69-247aa; ΔRecA1, Δ13-472aa; ΔRecA2, Δ493-681aa). **G**, Western immunoblot of V5, DLST and OGDH before and after immunoprecipitation using antibodies directed against V5, DLST, OGDH or non-specific immunoglobulins (IgG) in IMR5/75 cells expressing DDX1-V5 compared to DDX1-ΔSPRY(core), ΔRecA1 or ΔRecA2 truncation mutants, respectively. **H**, Western immunoblot of V5, DLST and OGDH before and after immunoprecipitation using antibodies directed against V5, DLST, OGDH or non-specific immunoglobulins (IgG) in IMR5/75 cells expressing DDX1-V5 or DDX1-V5-Δ269-295aa.I **I**, Representative confocal fluorescence imaging photomicrographs of proximity ligation assay signals (magenta dots) in IMR5/75 cells expressing DDX1-V5 or DDX1-V5-Δ269-295aa detected using anti-V5 and anti-DLST antibodies and counterstained with DAPI (blue) and phalloidin (yellow; scale bar: 7µm). **J**, Quantification of proximity ligation signal (magenta) using anti-DLST and anti-V5 antibodies in IMR5/75 cells expressing DDX1-V5 (*N* = 36) or DDX1-V5-Δ269-295aa (*N* = 34) as shown in Figure 5I. (Welch t-test, *P* = 1.476e-07). **K**, Western immunoblot of V5, P70-S6K, P70-S6K Thr389 phosphorylation and α-tubulin in KELLY cells after inducible expression of DDX1-V5, DDX1-V5-Δ269-295aa or an empty vector.

DDX ATPases usually contain a structurally conserved core with two RecA-like domains, which catalyze the enzymatic function of DDX proteins. Compared to other DDX ATPases, DDX1 has a unique protein structure. The first RecA-like domain is interrupted by a large SPla and the ryanodine receptor (SPRY) domain (Godbout et al., 1994), which is suspected to function as a protein-protein interaction platform (Kellner et al., 2015). To identify the protein domain required for DDX1:DLST interaction, we generated cells expressing a series of DDX1 domain truncation mutants tagged with V5 (Figure 5F). Truncation of the entire RecA-like domain 1, including the SPRY domain, strongly compromised the interaction with DLST, as evidenced by reduced co-immunoprecipitation (Figure 5G and Figure S6A). Conversely, no changes in DLST association were observed for a DDX1 truncation mutant lacking the RecA-like domain 2, hinting at the SPRY domain within RecA1 as a possible interaction site with DLST (Figure 5G). To test this, we generated a DDX1 mutant lacking the most conserved part of the SPRY domain (AA70-247), which contains the highly conserved surface patch predicted to serve as a protein interaction site (Figure 5F)(Kellner and Meinhart, 2015). Surprisingly, this DDX1 truncation mutant also preserved the interaction with DLST (Figure 5G), suggesting that the DDX1:DLST interaction may depend on the less conserved C-terminal part of the SPRY domain (AA247-295)(Kellner and Meinhart, 2015).

Disordered domains of proteins are candidate sites of protein-protein interaction (Hibino and Hoshino, 2020; Wong et al., 2020). To test whether a C-terminal part of the SPRY domain may provide the structural scaffold for the interaction with DLST, we searched for intrinsic disordered domains of DDX1, as predicted based on its amino acid sequences via PONDR (Xue et al., 2010) and IUPred2A (Mészáros et al., 2018). Polynomial modeling of the predicted interaction scores generated by these algorithms nominated amino acids 269aa to 295aa in DDX1 as a candidate disordered domain (Figure 5F). Indeed, immunoprecipitation of the V5-DDX1 Δ269-295aa mutant was associated with reduced DLST co-immunoprecipitation (Figure 5H). A proximality ligation assay (PLA) with V5-DDX1 Δ269-295aa-expressing cells confirmed the reduced interaction with DLST (Figure 5I and Figure 5J). This indicates that the amino acid stretch 269-295 in the C-terminal part of the SPRY domain of DDX1 is necessary for its interaction with DLST and raises the question whether this interaction is required for DDX1-mediated mTORC1 activation. Thus, DDX1 interacts with the α-KGDH complex in neuroblastoma cells, particularly when expressed at supraphysiological levels in the context of *DDX1-MYCN* co-amplification.

### DDX1:DLST interaction is required for DDX1-mediated mTORC1 pathway activation

As a key component of the α-KGDH complex, DLST plays an important role in the energy metabolism of cells. Since changes in energy metabolism also directly modulate mTORC1 pathway activation (de la Cruz López et al., 2019), we hypothesized that the interaction of DDX1 with DLST may interfere with α-KGDH function thereby stimulating mTORC1 activity. To assess whether the DDX1:DLST interaction was required for DDX1-mediated mTORC1 pathway activation, we ectopically overexpressed DDX1 or DDX1 Δ269-295aa in *MYCN*-amplified neuroblastoma cells. DDX1 Δ269-295aa was expressed at similar levels compared to wild-type DDX1, as confirmed by Western immunoblotting, and also localized to mitochondria (Figure S6B and S6C). Ectopic expression of DDX1 Δ269-295aa, however, was not associated with increased phosphorylation of P70-S6K at Thr389 in neuroblastoma cells (Figure 5K and Figure S6D), suggesting that it was insufficient to induce mTORC1 pathway activation. Thus, DDX1:DLST interaction is required for DDX1-mediated mTORC1 pathway activation.

### DDX1 alters *α*-KGDH complex activity resulting in *α*-KG accumulation and reduced oxidative phosphorylation (OXPHOS)

Considering the importance of DLST for α-KGDH complex activity in the TCA cycle (Tretter and Adam-Vizi, 2005), we hypothesized that the interaction between DDX1 and DLST may alter the catalytic function of α-KGDH and disrupt TCA cycle flux, which could lead to the accumulation of α-KG and subsequently activate mTORC1 to sustain cell survival, as previously reported in different contexts (Rathore et al., 2021). To test this hypothesis, we analyzed metabolomics data from the CCLE database including measurements of 225 metabolite levels in 928 cell lines from more than 20 cancer types (Li et al., 2019), and compared metabolite levels between cells with *DDX1-MYCN* co-amplification to those only harboring *MYCN* amplification. In line with altered α-KGDH activity, cancer cells with *DDX1-MYCN* co-amplification had higher levels of citrate, isocitrate, and α-KG (Figure 6A, Figure S7A and Table S5). Next, we measured metabolites in neuroblastoma cells ectopically expressing DDX1 compared to cells expressing the truncated DDX1 Δ269-295aa variant or cells expressing physiological levels of DDX1 performed using gas chromatography-mass spectrometry (GC-MS). Consistent with decreased α-KGDH activity, α-KG was increased upon ectopic DDX1 expression, but not when expressing DDX1 Δ269-295aa (Figure 6B). This indicates that aberrant DDX1 expression is sufficient to alter α-KG levels as a direct result of its interaction with the α-KGDH complex member DLST.

**Figure 6.**
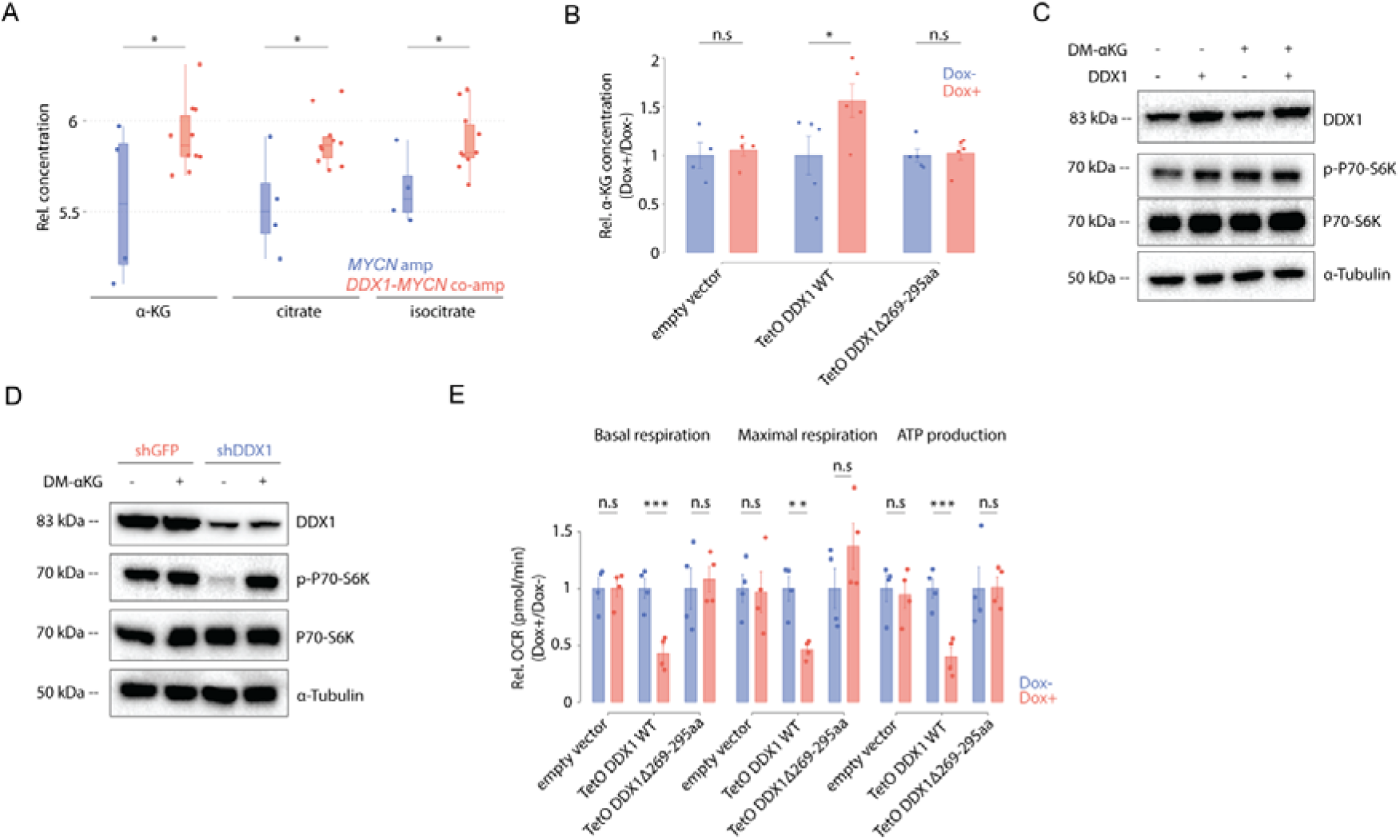
DDX1 hijacks the α-KGDH complex resulting in α-KG accumulation and OXPHOS reduction. **A**, Relative concentrations of α-KG, citrate and isocitrate in cancer cell lines with *DDX1*-*MYCN* co-amplifications (red) compared to cells only harboring *MYCN* amplifications (blue; Welch t-test, *P* = 0.038764, 0.008224 and 0.025814 for α-KG, citrate and isocitrate, respectively; *N* = 4 independent *MYCN*-amplified cancer cell lines versus *N* = 8 independent cancer cell lines with *DDX1*-*MYCN* co-amplification). **B**, Relative α-KG concentrations measured by GC-MS in KELLY cells ectopically expressing DDX1 or the DDX1-Δ269-295aa for 48 hours. KELLY cell transduced with an empty vector and exposed to doxycycline were used as control (Wilcox test, *P* = 0.02778; Data are shown as mean ± standard error) **C**, Western immunoblot of DDX1, P70-S6K, P70-S6K Thr389 phosphorylation and α-tubulin on KELLY cells treated with DM-αKG (2mM for 48 hours) in the presence or absence of ectopic DDX1 expression. **D**, Western immunoblot of DDX1, P70-S6K, P70-S6K Thr389 phosphorylation and α-tubulin in IMR5/75 cells treated with DM-αKG (2 mM for 48 hours) and expressing shRNA targeting either DLST or GFP as control. **E**, Mitochondrial oxygen consumption rate (OCR) measured using live-cell metabolic analysis at basal respiration, maximal respiration and ATP production in KELLY cells inducibly expressing DDX1 or DDX1-Δ269-295aa for 48 hours. KELLY cell transduced with a doxycycline-inducible empty vector served as negative control (Welch t-test, *P* = 0.002463, 0.01083 and 0.002127 for basal respiration, maximal respiration and ATP production, respectively; Data are shown as mean ± s.e.; *N* = 4 independent replicates).

As a rate-determining intermediate in the TCA cycle and the central product of glutaminolysis driving anaplerotic reactions in cells (Durán et al., 2012), α-KG can alter mTORC1 activity (Altman et al., 2016; Bodineau et al., 2021; de la Cruz López *et al*., 2019; Durán *et al*., 2012; Ge et al., 2018; Sancak et al., 2008; Takahara et al., 2020). We hypothesized that DDX1-mediated mTORC1 activation was due to α-KG accumulation. Indeed, incubation of neuroblastoma cell lines in the presence of membrane-permeable Dimethyl 2-oxoglutarate (DM-KG) was accompanied by mTORC1 pathway activation, phenocopying the effects of DDX1 overexpression (Figure 6C, 6D and Figure S7B). Disruption of α-KGDH is predicted to impair ATP production through OXPHOS in mitochondria. To test this, we measured oxygen consumption rates as a parameter to study mitochondrial function. Ectopic expression of DDX1, but not of the DDX1 Δ269-295aa mutant, was accompanied by reduced ATP production and respiration (Figure 6E, Figure S5F and Table S6). Thus, the DDX1:DLST interaction in cells expressing high levels of DDX1 can alter α-KGDH complex activity, resulting in α-KG accumulation, reduced OXPHOS and compensatory activation of mTORC1 pathway, which may explain the pronounced genetic dependency of *DDX1-MYCN* co-amplified cells on mTORC1.

### Pharmacologic mTORC1 inhibition results in cell death in cells with DDX1-MYCN co-amplification

Even though pharmacological mTOR inhibitors, such as everolimus and rapamycin, are in clinical use in patients suffering from different cancers, including *MYCN*-amplified neuroblastoma (Hua et al., 2019; Zou et al., 2020), biomarkers predicting mTOR inhibitor sensitivity are largely lacking. Besides its central role in TCA metabolism, α-KG can broadly influence cellular physiology, for example as the rate-limiting substrate of 2-oxogluterate-dependent dioxygenases in the management of hypoxia and in epigenetic remodeling (Liu et al., 2017; Losman et al., 2020; Wise David et al., 2011). Furthermore, α-KG accumulation can induce cancer cell differentiation and death (Abla et al., 2020; Morris et al., 2019; Zhang et al., 2021). Thus, we hypothesized that the α-KG-induced activation of the mTORC1 pathway in cells with *DDX1-MYCN* co-amplification was required to sustain cancer cell viability through mTORC1-dependent cell survival mechanisms (Durán *et al*., 2012; Ferrara-Romeo et al., 2020; Mills et al., 2008). To test this hypothesis, we incubated *MYCN*-amplified neuroblastoma cell lines with and without *DDX1* co-amplification with DM-αKG in the presence and absence of mTORC1 inhibitors. Indeed, cells were more sensitive to DM-αKG induced cell death when overexpressing DDX1, an effect that was potentiated when combined with the mTORC1 inhibitor rapamycin (Figure S8A-D). Moreover, cell lines harboring a *DDX1*-*MYCN* co-amplification were more sensitive to rapamycin treatment than cell lines only harboring a *MYCN* amplification (Figure 7A and Figure S8E). In line with DDX1-induced mTORC1 dependency, ectopic expression of DDX1 increased sensitivity to rapamycin, which was not observed when expressing mutant DDX1 Δ269-295aa (Figure 7B and Figure S8F), suggesting that the increase in sensitivity depended on the DDX1-DLST interaction. In turn, shRNA-mediated DDX1 knockdown in neuroblastoma cells with a *DDX1*-*MYCN* co-amplification resulted in reduced rapamycin sensitivity (Figure 7C and Figure S8G), indicating that high DDX1 expression was required for mTOR inhibitor sensitivity in the context of *DDX1*-*MYCN* co-amplification. Previously published 50% inhibitory concentrations (IC_50_) for rapamycin from the Genomics of Drug Sensitivity in Cancer database (GDSC2)(Iorio et al., 2016; Yang et al., 2013) anti-correlated significantly with *DDX1* copy number in *MYCN*-amplified neuroblastoma cell lines (Figure 7D, Pearson coefficient = −0.5043 in neuroblastoma cells, *P* = 0.0394), further corroborating the link between *DDX1* co-amplification and mTORC1 dependency. Lastly, we treated primary zebrafish neuroblastic tumor cells either co-expressing DDX1 and MYCN or MYCN alone with rapamycin (Figure S8H). DDX1-MYCN expressing neuroblastic zebrafish cells were more sensitive to pharmacological mTORC1 inhibition than cells only expressing MYCN (Figure 7E). In conclusion, high DDX1 expression as a result of *DDX1*-*MYCN* co-amplification can induce a therapeutically actionable dependency on mTORC1.

**Figure 7.**
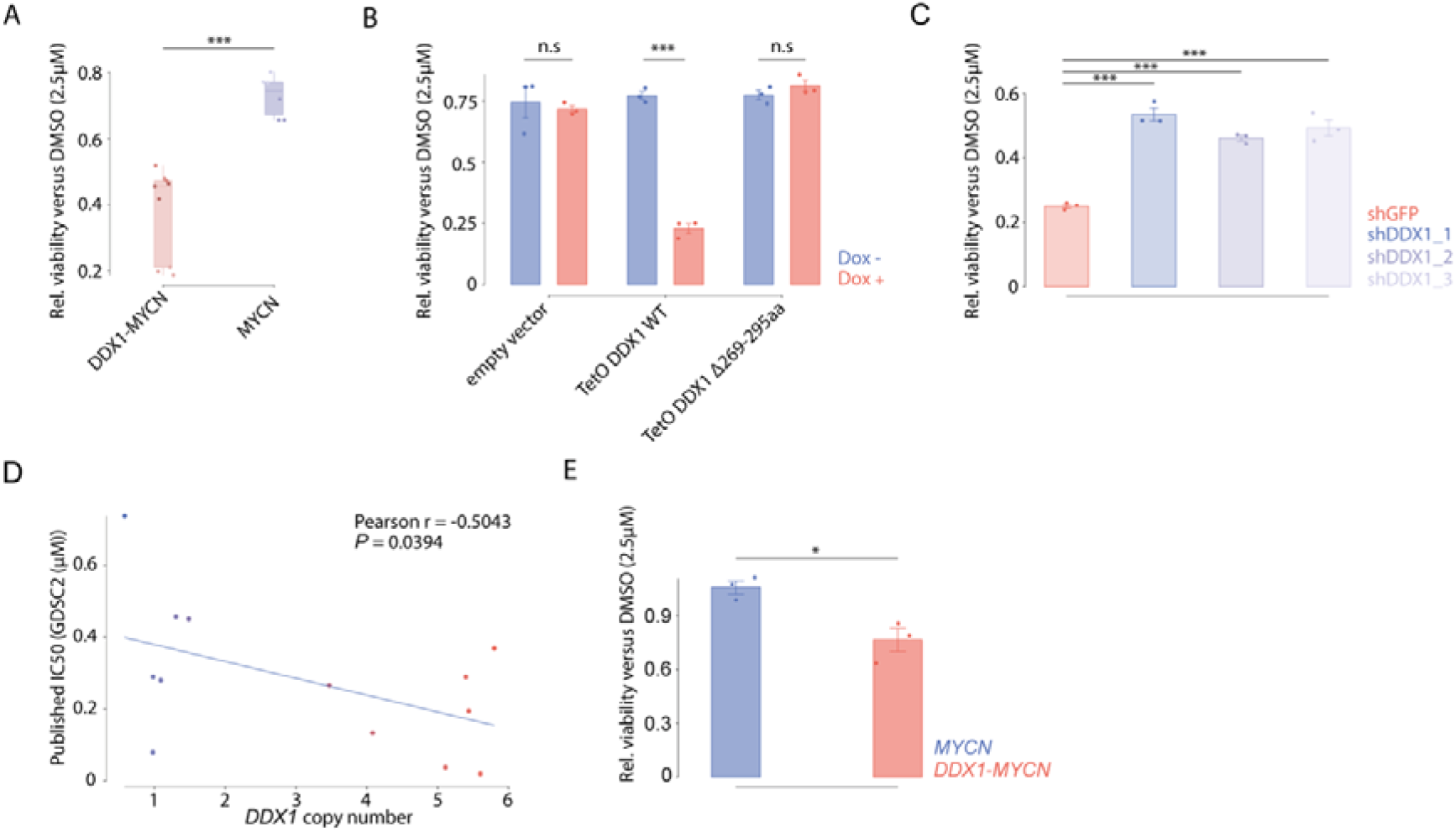
Aberrant DDX1 expression is sufficient to increase sensitivity to pharmacological mTORC1 inhibition. **A**, Relative cell viability as measured using MTT assay of neuroblastoma cell lines with *DDX1*-*MYCN* co-amplification (red, *N*= 3), or with *MYCN* amplifications (blue, *N* = 2) treated with rapamycin (2.5 µM for 72 hours) compared to cell viability after DMSO (vehicle control) treatment (Welch t-test, *P* = 2.291e-05 *DDX1*-*MYCN* vs *MYCN*; each data point represents a technical replicate). **B**, Relative cell viability as measured using MTT assay of KELLY cells inducibly expressing DDX1, DDX1-Δ269-295aa or an empty vector and treated with rapamycin (2.5 µM for 72 hours) compared to cell viability after DMSO (vehicle control) treatment (Welch t-test, *P* = 3.943e-05; data are shown as mean ± s.e.; *N* = 3 technical replicates). **C**, Relative cell viability as measured using MTT assay of IMR5/75 cells expressing shRNAs directed against DDX1 (blue) or GFP (red) and treated with rapamycin (2.5 µM for 72 hours) compared to cell viability after DMSO (vehicle control) treatment. (Pairwise t-test adjusted by Benjamini-Hochberg correction, *P* = 1.3e-05, 4.1e-05 and 2.2e-05 for each independent shRNA directed against DDX1 vs. shGFP, respectively; Data are shown as mean ± s.e.; *N* = 3 technical replicates). **D**, Correlation between the *DDX1* copy number and the IC_50_ value of rapamycin in different neuroblastoma cell lines derived from the GDSC2 database (Pearson correlation, *R* = −.05043, *P* = 0.0394, *N* = 13 independent cancer cell lines). **E**, Relative cell viability as measured using MTT assay of neuroblastic tumor cells derived from transgenic zebrafish expressing MYCN or MYCN and DDX1 and treated with rapamycin (2.5 µM for 72 hours) compared to cell viability after DMSO (vehicle control) treatment (Welch t-test, *P* = 0.02707; Data are shown as mean ± s.e.; *N* = 3 independent replicates from cells derived from different zebrafish).

## Discussion

With the goal to expand therapeutic strategies in cancer beyond targetable molecular alterations, we found that the co-amplification of a passenger gene, which is not directly involved in tumorigenesis, can create pharmacologically actionable amplicon structure-defined collateral lethal therapeutic vulnerabilities. Our re-analysis of pan-cancer genomes further suggests that this strategy may be successful in many cancer entities with diverse DNA amplicons.

We and others have previously shown that large neighboring genomic regions harboring enhancers co-amplify with oncogenes on the same intra- or extrachromosomal DNA amplicon (ecDNA)(Helmsauer *et al*., 2020; Morton *et al*., 2019). This implies that positive selection acts on these rewired loci. Here, we describe that the co-amplification of neighboring genomic regions frequently also results in the inclusion of passenger genes. Under our current model, passenger genes are under neutral selection and represent mere structural bystanders of DNA amplifications. However, it is conceivable that some passenger genes on amplicons provide functional or structural advantages to cancer cells. Functionally, passenger genes could improve tumor cell fitness under special cellular or environmental conditions. Structurally, yet unidentified elements near or within passenger genes may also positively influence ecDNA stability, maintenance or oncogene regulation. Thus, we here provide an additional layer of information on the content of DNA amplicons, which may help resolve longstanding questions about their structural requirements and functional consequences.

Some new important questions for cancer therapy and treatment resistance directly arise from our observations: Firstly, there are other passenger genes on the *MYCN* amplicon, e.g., *NBAS* and *FAM49A*, which might similarly create their own, so far unidentified therapeutic vulnerabilities. Identifying such collateral vulnerabilities may allow new therapeutic approaches that would substantially improve tumor eradication in high-risk MYCN-driven cancers. Beyond the idea of targeting individual vulnerabilities created by the co-amplification of different passenger genes, the *MYCN* amplicon also contains a unique chromatin landscape with enhancers required to drive gene expression from the amplicon (Helmsauer *et al*., 2020; Morton *et al*., 2019). It is tempting to speculate that the structural coupling of genes and their coordinated expression from the joint enhancers could create additional, amplicon-specific, therapeutically actionable vulnerabilities.

The DEAD-box ATPase DDX1 has previously been implicated in various steps of DNA, mRNA, rRNA and tRNA processing and repair. Our in-depth investigation of *DDX1* co-amplification revealed a previously unanticipated and fundamentally new role of DDX1 in cellular metabolism by uncovering its interaction with the α-KGDH complex as a non-canonical interaction partner of the DLST subunit in neuroblastoma cells. Our data suggest that high DDX1 expression impedes the TCA cycle as well as OXPHOS and consequently promotes accumulation of α-KG, which in turn triggers mTORC1 activation to maintain tumor cell survival. Many nuclear ATPases localize to mitochondria (Ding and Liu, 2015; Padmanabhan et al., 2016), but to our knowledge interactions of these ATPases with TCA enzymes or a direct impact on cellular metabolism have not been reported to date. DDX1 is unique amongst other DDX protein family members, because it contains a long ∼211 amino-acid insertion between the signature motifs of their ATPase core (Godbout *et al*., 1994). This insertion encompasses the SPRY core domain and at its C-terminal end a relatively unconserved presumably disordered domain, which we found to be essential for the interaction between DDX1 and DLST. Additionally, DDX1 is functionally unique in its exceptionally high affinity for ADP, which could leave DDX1 in an inactive, ADP bound form at cytoplasmic nucleotide concentrations. Whether the spatial enrichment of ATP in mitochondria facilitates conformational changes of DDX1 and thereby promotes substrate exchange or whether so far unidentified nucleotide exchange factors exist to promote ADP release remains to be determined. It also needs to be tested if DDX1 interacts with DLST at physiological protein levels in untransformed cells, or if this only occurs in the context of aberrant DDX1 expression in *DDX1-MYCN-*amplified cancer cells. Since the ATPase activity of DDX proteins are required for their role in DNA and RNA biology, it is conceivable that these functions are not required for DDX1 to establish stable interactions with protein substrates such as a-α-KGDH or as recently reported with casein kinase 2 (Fatti et al., 2021). Whether such canonical DDX1 functions also influence mitochondrial metabolism and thereby contribute to the observed mTORC1 dependency remains to be investigated.

Recent reports suggest that the α-KGDH complex can regulate histone succinylation and gene expression in the nucleus (Wang et al., 2017). DDX1 and its DNA:RNA binding has important functions in many cell activities also located in the cell nucleus (Chen *et al*., 2002; Han *et al*., 2014; Li *et al*., 2008; Zhang *et al*., 2011). Therefore, DDX1 could in principle also interact with DLST in the nucleus and may affect the role of α-KGDH in histone succinylation and gene expression, which could also generate, yet undefined, collateral vulnerabilities or may contribute to the observed mTORC1 dependency.

Even though mTORC1 is a well-studied multiprotein complex essential for cancer cell survival, proliferation, and growth (Guertin and Sabatini, 2007; Kim et al., 2017; Mills *et al*., 2008), biomarkers predicting patient responses to mTOR inhibitors are still largely missing. Intriguingly, rapamycin is part of the treatment protocol RIST (rapamycin, irinotecan, sunitinib, temozolomide), which was developed for treatment-refractory or relapsed neuroblastomas (Corbacioglu et al., 2013). Even though rapamycin is only one of four agents used in this treatment protocol, it will be important to test whether neuroblastoma patients with *DDX1-MYCN* amplifications respond better to RIST than neuroblastoma patients without such co-amplifications. These analyses may reveal that *DDX1-*co-amplificaiton could serve as a predictive response biomarker for mTORC1 inhibitor treatment.

We and others recently identified ecDNA copy number dynamics as a driver of therapy resistance (Lange et al., 2021). Based on these observations, we anticipate that decreases in the number of *DDX1*-containing ecDNA under mTORC1 inhibitor treatment could represent one way of treatment evasion in *DDX1-MYCN*-amplified cancer cells. The concept of ecDNA-targeting therapies has recently been proposed (van Leen *et al*., 2022). Since such therapies may prevent ecDNA-mediated resistance acquisition, combining targeted therapies based on amplicon-structure-defined collateral vulnerabilities with ecDNA-directed therapies may represent therapeutically meaningful future avenues.

In conclusion, we here present a strategy to eliminate cancer cells by targeting factors not directly linked to cancer pathogenesis. We identified a collateral vulnerability in neuroblastoma cells, which is created through passenger-mediated metabolic reprogramming. We propose that pharmacological mTORC1 inhibition could provide an effective therapy for a meaningful fraction of cancer patients with *DDX1*-*MYCN* co-amplification. Since passenger co-amplifications are common in cancer, our approach has the potential to identify previously unanticipated therapeutic targets and transform target discovery in oncology, especially in cancers with amplification of oncogenes that have been considered undruggable to date.

## STAR⍰Methods

### Cell culture

Human tumor cell lines were obtained from the American Type Culture Collection (ATCC) or a gift from collaborative laboratories. The identity of all cell lines was verified by short tandem repeat genotyping (Eurofins Genomics). The absence of *Mycoplasma* sp. contamination was determined using a Lonza MycoAlert system (Lonza). Neuroblastoma cell lines were cultured in RPMI-1640 medium (Gibco) supplemented with 1 % of penicillin, streptomycin, and 10 % of fetal calf serum (FCS) (Thermo Fisher). RPE cells were cultured in DMEM (Gibco) supplemented with 1 % of penicillin, streptomycin, and 10 % of FCS. To assess the number of viable cells, cells were trypsinized (Thermo Fisher), resuspended in medium, and sedimented at 300 *g* for 5 minutes. Cells were then resuspended in medium, mixed in a 1:1 ratio with 0.02 % trypan blue (Thermo Fisher), and counted with a Bio-Rad TC20 cell counter.

### Cell viability measurements

10,000 cells per well were seeded in transparent, flat-bottom, 96 well plates. After 24 hours, drug was added to the medium and cells were incubated for 72 hours. 3-(4,5-dimethylthiazol-2-yl)-2,5-diphenyltetrazolium (MTT) assay reagent (Abcam, ab211091) was added according to the manufacturers protocol, and MTT signal was measured by an Epoch plate reader (BioTeK) with read absorbance at OD = 590nm.

### Plasmid constructs

Human *DDX1* cDNA (NM_004939.2) was PCR-amplified and isolated from pRecLV151-DDX1 (GeneCopoeia, Rockville, MD, USA). DDX1 cDNA was cloned into pENTR1A (Thermo Fisher) using restriction enzymes SalI and NotI (New England Biolabs) and cloned into a pInducer20 (Addgene) using the Gateway strategy and the manufacturer’s protocol (Thermo Fisher). DDX1 cDNA was cloned into the pRNTR1A vector in frame with C-terminal V5 tag or mCherry and used to generated pInducer20-DDX1-V5 or pInducer20-DDX1-mcherry using the Gateway cloning, according to the manufacturer’s instructions. Large truncation of V5-tagged DDX1 lentiviral vectors were generated using site-directed mutagenesis according to the manufacturer’s instructions (Q5® Site-Directed Mutagenesis, New England Biolabs) and were confirmed using sanger sequencing. Truncation of core SPRY domain in DDX1: Truncation of unordered region of SPRY domain in DDX1: K69-F247 is missing. Truncation of RecA-like domain 1 in DDX1: I13-K472 is missing. Truncation of RecA-like domain 2 in DDX1: K493-V681 is missing. Human *DLST* cDNA (NM_001933.4) was PCR-amplified and isolated from human retro-synthesized cDNA library. DLST cDNA was cloned into pENTR1A using restriction enzymes SalI and NotI (New England Biolabs) and cloned into a pInducer20 using the Gateway strategy and the manufacturer’s protocol (Thermo Fisher). pLKO.1 shRNA plasmids targeting DDX1 (TRCN0000050500, TRCN0000050501, TRCN0000050502) and control targeting GFP (shGFP) were obtaining from the RNAi Consortium (Broad Institute).

### Lentivirus production and cell transduction

Lentivirus production was carried out as previously described (Henssen et al., 2017). In brief, HEK293T cells were transfected with TransIT-LT1 (Mirus) in a 2:1:1 ratio of the lentiviral vector and psPAX2 and pMD2.G packaging plasmids (Addgene), according to the manufacturer’s instructions. Viral supernatant was collected 48 and 72 hours after transfection. The supernatant was pooled, filtered, and stored at –80°C. Neuroblastoma and RPE cells were transduced with virus particles in the presence of 8 μg/ml hexadimethrine bromide (Merck). Cells were transduced for 1 day in antibiotic-free medium and then grown in full medium for 1 day. Neuroblastoma cells were then selected for 2 days with puromycin hydrochloride (2 μg/ml) or geneticin disulphate (G418, Roth) (2mg/ml).

### Western immunoblotting

Whole-cell protein lysates were prepared by lysing cells in 15 mM HEPES, 150 mM NaCl, 10 mM EGTA, and 2 % (v/v) Triton X-100 supplemented with cOmplete (Roche) and PhosStop (Roche) phosphatase inhibitors. Protein concentrations were assessed by bicinchoninic acid assay (BSA, Santa Cruz Biotechnology). For 5 minutes, 10 μg of protein was denatured in Laemmli buffer at 90 °C. Samples were run on NuPage 10 % polyacrylamide, 1 mm Tris-Glycin Protein Gels (Thermo Fisher Scientific) and transferred to PVDF membranes (Roche). Membranes were blocked with 5 % dry milk or 5 % BSA (Roth) in TBS with 0.1 % (v/v) Tween-20 (Carl Roth). Membranes were probed with primary antibodies overnight at 4 °C and then with secondary antibodies conjugated to horseradish peroxidase for 1 hour at room temperature. Chemiluminescent detection of proteins was carried out using Immunocruz Western blotting luminol reagent (Santa Cruz Biotechnology) and the Fusion FX7 imaging system (Vilber Lourmat). Densitometry was performed using ImageJ (NIH).

### Immunofluorescence staining and colocalization analysis

Cells were grown at the desired confluence on a glass cover slide for 24 hours and treated with 1000ng/mL doxycycline for another 48 hours (for the corresponding experiment). Cells were washed with phosphate-buffered saline (PBS) three times and fixed for 10 minutes with 4 % paraformaldehyde, washed with PBS three times and permeabilized with PBS containing 0.2 % Triton-X100. For immunofluorescence, cells were blocked for 30 minutes in 10 % FCS in PBS, incubated overnight at 4ºC with the primary antibody, washed three times with PBS-T (0.05 % Tween-20 in PBS), incubated for 1 hour in the dark at room temperature with the secondary antibody, washed three times with PBS-T and mounted on a slide with 4′,6-diamidino-2-phenylindole (DAPI)-containing mounting media. As co-localization staining, DDX1-mCherry or DDX1-mCherry-Δ269-295aa inducibly expressed KELLY cells were seeded on 8 well µ-slide (ibidi) for 48 hours in the presence or absence of doxycycline (1µg/ml, Sigma-Aldrich). 30 minutes before fixation, cells were incubated with MitoTracker (500nM, cell signaling technology) and Hoechest (1µg/ml, Thermo Fisher) at 37ºC. After fixation, cells were washed 3 times with PBS and mounted with PBS. Cells were imaged using a Leica TCS SP5 II (Leica Microsystems) and quantified using ImageJ.

### RNA sequencing

mRNA was isolated from DDX1 inducibly expressed KELLY cells after 48 hours incubation in the presence or absence of doxycycline (1µg/ml, Sigma-Aldrich). Libraries were sequenced on HiSeq 2000 v4 instruments with 2×125-bp paired-end reads (Illumina). Reads were mapped with STAR(v2.7.6a) to the human reference genome hg19 with the Gencode v19 annotation using default parameters (Dobin et al., 2013). Gene abundance was estimated using RSEM (v1.3.1) (Li and Dewey, 2011), counting only alignments with both mates mapped and allowing for fractional counting of multi-mapping and multi-overlapping reads.

### Clonogenic assay

5,000 cells were seeded in 24 well plate coated with poly-l-lysin (Merck, USA). After 24 hours, drugs and doxycycline (1µg/ml) were added to the medium and fresh medium with drugs and doxycycline was replaced every 48 hours. Cells were continuously cultured for 10 days until formation of colonies was observed. Cells were fixed with 3.7 % formaldehyde for 10 minutes at room temperature, dried and stained with 0.1 % crystal violet (Merck) in 10 % ethanol (Roth) for 10 minutes. After washing with sterile water and drying, colonies were measured by ColonyArea (Guzmán et al., 2014) from ImageJ (Schneider et al., 2012).

### Proximity ligation assay

Cells were seeded into 8 well slides at 3000 cells per well, treated for 48 hours with doxycycline to induce V5-DDX1 and V5-DDX1(Δ269-295) expression. After fixation for 10 minutes with 4 % paraformaldehyde and blocking for 30 min with 10 % FCS in PBS, cells were incubated overnight at 4ºC with primary antibody against V5 (Mouse, Abcam, 1:500) and DLST (Rabbit, Cell Signaling Technology, 1:500). Proximity ligation assay was performed using Duolink® In Situ Kit (Sigma-Aldrich) according to the manufacturer’s protocol. Nuclei were counterstained using Duolink® In Situ Mounting Medium with DAPI (Sigma-Aldrich) and F-actin were stained using Phalloidin (Thermo Fisher) according to the manufacturer’s instruction. Pictures were taken with a Leica TCS SP5 II (Leica Microsystems) with 63-fold magnification and analyzed using ImageJ.

### Zebrafish maintenance

Zebrafish (*Danio rerio*) were raised and maintained according to standard protocols at 28°C with a 14/10h light dark cycle (Westerfield, 2000). The transgenic line Tg (*dβh*-*MYCN*: *dβh*-*eGFP*) (Tao et al., 2017) was a kind gift from Thomas Look (Dana Faber Cancer Institute, Boston, USA). All zebrafish were of the AB background strain. All experiments were performed in accord with the legal authorities approved license “G 0325/19”.

### Zebrafish transgenesis

Zebrafish line Tg (*dβh*-*MYCN*:*dβh*-*eGFP*) was a kind gift of Thomas Look (Dana Faber Cancer Institute, Boston, USA) and described previously (Tao et al., 2017). Plasmid *dβh-eGFP (pDest_IsceI)* was also a kind gift of Thomas Look. Plasmid of Tol2 constructs were a kind gift from Jan Philipp Junker. To create *dβh-DDX1-polyA-Tol2-CryAA:mCerulean*, the *dβh* promoter was excised using restriction enzymes ClaI and KpnI (New England Biolabs) and cloned in p5E-MCS (Multiple Cloning Sites, Addgene #26029), linearized with the same enzymes, through T4 Rapid DNA Ligation Kit (Roche). DDX1 was PCR-amplified using primers containing suitable recombination sites for the Gateway System (FW: GGGGACAAGTTTGTACAAAAAAGCAGGCTTACCATGGCGGCCTTCT, RV: GGGGACCACTTTGTACAAGAAAGCTGGGTTCTAGAAGGTTCTGAACAGCTGGTT AGG). DDX1 fragment was cloned into pDONR221 (Thermo Fisher) using the Gateway system following the manufacturer’s protocol (Thermo Fisher). P3E-polyA and pDEST-Tol2-*CryAA-mCerulean* were a kind gift of Jan-Philipp Junker (Max Delbrück Center, Berlin, Germany). P5E-*dβh*, pDONR-DDX1 and p3E-polyA were cloned in a pDEST-Tol2-*CryAA*-*mCerulean* using the Gateway System following the manufacturer’s protocol (Thermo Fisher). The final construct was sequenced by LightRun Sequencing (Eurofins Genomics) to confirm the successful reaction. To generate the zebrafish transgenic line Tg (*dβh*-*DDX1*:*CryAA*-*mCerulean*), plasmid *dβh*-DDX1-polyA-Tol2-*CryAA*:*mCerulean* was injected into fertilized eggs and fish were grown to adulthood.

### Zebrafish tumor cell treatment

Tumors from Tg (*dβh*-*MYCN*:*dβh* -*eGFP*; *dβh*-*DDX1*:*CryAA*-*mCerulean*) double transgenic fish and Tg (*dβh*-*MYCN*:*dβh*-*eGFP*) were excised from adult fish immediately after hypothermal shock euthanasia. Tumors were dissociated using a Collagenase II (Thermo Fisher) based protocol (330 µL Collagenase II – final concentration: 100 U/mL -, 120 µL HBSS, 50 µL FCS for 30 minutes at 37 °C, followed by 5 minutes of incubation at 37 °C after addition of 200 µL of Dispase II (Thermo Fisher) – final concentration: 2 U/mL -, gently pipetting every 10 minutes). After dissociation, single cell suspension was transferred to Round-Bottom Polystyrene Test Tubes with Cell Strainer Snap Cap to remove undissociated tissue. Cells were resuspended in DMEM (Gibco) supplemented with 10 % FCS and 1 % Penicillin/Streptomycin and plated in 96 well plates at a density of 0,1 × 10^5^ cells. Cells were incubated at 27 °C, 5 % CO2. After 24 h, cells were visually inspected for eGFP expression using a AXIO microscope (Zeiss) and Rapamycin (2.5 µM) or vehicle were added to the DMSO; cells were then incubated for 72 h and viability assessed through MTT assay (Abcam) following manufacturer’s protocol. Absorbance was measured using an Epoch plate reader (BioTeK) at OD = 590.

### Oxygen consumption rate (OCR) measurements

The mitochondrial respiratory capacity was determined with the XF Cell Mito Stress Test Kit (Agilent Technologies). Cells were seeded in the XF96 cell culture microplate at a density of 1 × 10^4^ per well with 4 replicates of each condition. XF96 FluxPak sensor cartridge was hydrated with Seahorse Calibrant overnight in a non-CO_2_ incubator at 37 °C. The following day, cells were incubated with the Seahorse medium (plus 1 mM pyruvate, 2 mM glutamine, and 10 mM glucose) for 1 hour prior. The OCR was measured by Xfe96 extracellular flux analyzer with the sequential injection of 1 μM oligomycin A, 0.5 μM carbonyl cyanide-p-trifluoromethoxyphenylhydrazon (FCCP), and 0.5 μM rotenone/antimycin A.

### Electron microscopy

Cells were grown on poly-l-lysin-coated sapphire discs and frozen using the Leica EM ICE. Freeze substitution was done in 1 % H_2_O (v/v), 1 % glutaraldehyde (v/v), 1 % osmium tetroxide (v/v) in anhydrous acetone using the following protocol: 37h at −90 °C, 8h from −90 to −50 °C, 6h from −50 to −30 °C, 12h at −20 °C and 3h from −20 to 20 °C. Samples were further contrasted with 0.1 % uranyl acetate [w/v] in anhydrous acetone and infiltrated with 30 %, 70 % and 90 % epon-acetone mixtures for 2h each, followed by 3 × 2h changes of 100 % epon (Polybed 812, Science Services) and polymerized at 60 °C for 48 h. 70nm sections were obtained with an ultra-microtome and imaged at 80 kV with an EM910 (Zeiss). ImageJ was used for quantification.

### Co-immunoprecipitation

In the standard co-immunoprecipitation assay, cells were lysed in lysis buffer (50□mM Tris pH□7.5, 150□mM NaCl, 10mM MgCl_2_, 0.5 % Nonidet P40 (Igepal), 10 % Glycerol, 1 mM NaF, freshly added 1□mM 4-(2-Aminoethyl)benzenesulfonyl fluoride hydrochloride (AEBSF, Sigma-Aldrich), protease inhibitors) and frozen in liquid nitrogen for 1□min. After thawing at 37 ºC shortly and removal of cell debris by centrifugation, 1.2□μg of antibody and 100 µl of 20 % Protein A-Sepharose beads (Amersham Biosciences) were added to clarified whole cell extract (WCE) and incubated overnight at 4□°C. The next day, beads then were subjected to three washes with lysis buffer contained 1mM Dichlorodiphenyltrichloroethane (DTT, Thermo Scientific). The beads were then sent for Mass Spectrometry based Proteomics of DDX1 interactome or boiled with 1× Sodium dodecyl sulfate (SDS) loading buffer at 90 ºC for western blot analysis. For V5-tagged immunoprecipitation, to assess the binding region of DDX1 to α-KGDH complex, different truncated V5-DDX1s were overexpressed. The same amounts of input and V5 immunoprecipitation eluates were loaded in western blot analysis for the detection of α-KGDH complex.

### Mass Spectrometry based Proteomics

Beads from immunoprecipitation experiments were resuspended in 20 mL denaturation buffer (6 M Urea, 2 M Thiourea, 10 mM HEPES, pH 8.0), reduced for 30 min at 25 °C in 12 mM dithiothreitol, followed by alkylation with 40 mM chloroacetamide for 20 min at 25 °C. Samples were first digested with 0.5 µg endopeptidase LysC (Wako, Osaka, Japan) for 4 h. After diluting the samples with 80 µl 50 mM ammonium bicarbonate (pH 8.5), 1 µg sequence grade trypsin (Promega) was added overnight at 25 °C. The peptide-containing supernatant was collected and acidified with formic acid (1 % final concentration) to stop the digestion. Peptides were desalted and cleaned up using Stage Tip protocol(Rappsilber et al., 2003). After elution with 80 % acetonitrile/0.1 % formic acid, samples were dried using speedvac, resolved in 3 % acetonitrile/0.1 % formic acid and analysed by LC-MS/MS.

Peptides were separated on a reversed-phase column (20 cm fritless silica microcolumns with an inner diameter of 75 μm, packed with ReproSil-Pur C18-AQ 1.9 μm resin (Dr. Maisch GmbH) using a 90 min gradient with a 250 nL/min flow rate of increasing Buffer B concentration (from 2 % to 60 %) on a High-Performance Liquid Chromatography (HPLC) system (Thermo Fisher) and ionized using an electrospray ionization (ESI) source (Thermo Fisher) and analyzed on a Thermo Q Exactive HF-X instrument. The instrument was run in data dependent mode selecting the top 20 most intense ions in the MS full scans, selecting ions from 350 to 2000 *m*/*z*, using 60 K resolution with a 3 × 10^6^ ion count target and 10 ms injection time. Tandem MS was performed at a resolution of 15 K. The MS2 ion count target was set to 1 × 10^5^ with a maximum injection time of 22 ms. Only precursors with charge state 2–6 were selected for MS2. The dynamic exclusion duration was set to 30 s with a 10-ppm tolerance around the selected precursor and its isotopes.

Raw data were analyzed using MaxQuant software package (v1.6.3.4)(Tyanova et al., 2016a). The internal Andromeda search engine was used to search MS2 spectra against a human UniProt database (HUMAN.2019-07) containing forward and reverse sequences. The search included variable modifications of methionine oxidation, N-terminal acetylation and fixed modification of carbamidomethyl cysteine. Minimal peptide length was set to seven amino acids and a maximum of 3 missed cleavages was allowed. The FDR was set to 1 % for peptide and protein identifications. Unique and razor peptides were considered for quantification. Retention times were recalibrated based on the built-in nonlinear time-rescaling algorithm. MS2 identifications were transferred between runs with the “Match between runs” option in which the maximal retention time window was set to 0.7 min. The LFQ (label-free quantitation) algorithm was activated.

The resulting text file was filtered to exclude reverse database hits, potential contaminants, and proteins only identified by site. Statistical data analysis was performed using Perseus software (v1.6.2.1)(Tyanova et al., 2016b). Log2 transformed LFQ intensity values were filtered for minimum of 3 valid values in at least one experimental group and missing values were imputed with random low intensity values taken from a normal distribution. Differences in protein abundance between DDX1 bait and IgG control samples were calculated using two-sample Student’s t-test. Proteins enriched in the DDX1 group and passing the significance cut-off (permutation-based FDR□<□5 %), were defined as DDX1 interactors.

### Gas chromatography–mass spectrometry (GS-MS)

Cells were lysed with 5 mL of ice-cold 50% methanol (MeOH, Honeywell) solution containing 2 µg/mL cinnamic acid (Sigma-Aldrich). Immediately after the MeOH solution was added to the culture plate, lysates were scraped into the MeOH solution and the methanolic lysates were collected. After cell harvest, 4 mL of chloroform (CHCl_3_, VWR), 1.5 mL of MeOH and 1.5 mL of water (H_2_O, VWR) was added to the methanolic cell extracts, shaken for 60 minutes at 4 °C, and centrifuged at 4,149xg for 10 minutes to separate the phases. The polar phase (6 mL) was collected and dried at 30 °C at a speed of 1,550x g at 0.1 mbar using a rotational vacuum concentrator (RVC 2-33 CD plus, Christ, Osterode am Harz, Germany). Samples were pooled after extraction and used as a quality control (QC) sample to test the technical variability of the instrument. They were prepared alongside the samples in the same way. After drying, samples were split by adding 600 µL of 20% MeOH to the dried extracts, shaking for 60 minutes at 4 °C, followed by centrifugation at maximum speed (18,213xg) for 10 minutes. Two 280 µL aliquots per sample were then dried under vacuum of which one was analyzed and the other kept as a backup.

All polar cell extracts were stored dry at −80 °C until analysis. Extracts were removed from the freezer and dried in a rotational vacuum concentrator for 60 minutes before further processing to ensure there was no residual water, which may influence derivatization efficiency. Dried extracts were dissolved in 15 µL of methoxyamine hydrochloride solution (40 mg/mL in pyridine) and incubated for 90 minutes at 30 °C with constant shaking, followed by the addition of 50 µL of N-methyl-N-[trimethylsilyl]trifluoroacetamide (MSTFA) including an alkane mixture for retention index determination and incubated at 37 °C for 60 minutes. The extracts were centrifuged for 10 minutes at 10,000 x g, and aliquots of 25 µL were transferred into glass vials for GC-MS measurement. An identification mixture for reliable compound identification was prepared and derivatized in the same way and an alkane mixture for reliable retention index calculation was included(Opialla et al., 2020).

Metabolite analysis was performed on a Pegasus BT GC-TOF-MS-System (LECO Corporation, St. Joseph, MN, USA) complemented with an auto-sampler (Gerstel). Gas chromatographic separation was performed on an Agilent 8890 (Agilent Technologies, Santa Clara, CA, USA), equipped with a VF-5 ms column of 30-m length, 250-μm inner diameter, and 0.25-μm film thickness (Agilent technologies, Santa Clara, CA, USA). Helium was used as carrier gas with a 1.2 mL/min flow rate. Gas chromatography was performed with the following temperature gradient: the first 2 minutes allowed the column to equilibrate at 70°C, a first temperature gradient was applied at a rate of increase of 5°C per minute until a maximum temperature of 120°C was reached. Subsequently, a second temperature gradient was applied using a rate of increase of 7°C/min up to a maximum temperature of 200°C. This was immediately followed by a third gradient of 12°C/min up to a maximum temperature of 320°C with a hold time of 7.5 min. The spectra were recorded in a mass range of 60 to 600 m/z with a scan rate of 10 spectra/s. A split ratio of 1:5 was used. QC samples were used both for conditioning (6 mouse liver samples at the beginning of the run) the instrument and for measuring technical variability (pooled QC samples) across the batch. The pooled QC samples were run at the beginning and end of each batch and after every 10th sample. The GC-MS chromatograms were processed with the ChromaTOF software (LECO Corporation, St. Joseph, MN, USA) including baseline assessment, peak picking, and computation of the area and height of peaks without calibration by using an in-house created reference and a library containing the top 3 masses by intensity for metabolites related to the central carbon metabolism. Data were normalized to the sum of the area. Individual derivatives were summed up. Relative quantities were used. The quality control samples were analyzed separately (Table S7).

### Copy number analysis

The Patients’ copy number dataset of Pan-Cancer Analysis of Whole Genomes (PCAWG) study and Tumor Alteration Relevant for Genomics-driven Therapy (TARGET) were retrieved from cbioportal database (https://www.cbioportal.org). The high-level amplified genes were labeled as “2” from the profile description. In copy-number data for 556 neuroblastoma patients, cutoffs were chosen to maximize discrimination between *MYCN*-amplified and nonamplified samples (or *DDX1*-amplified and nonamplified samples) and exclude high-level gains: 1.5 for Affymetrix and NimbleGen arrays, 2 for Agilent arrays and 0.7 for Illumina arrays. Finally, segments with log2 ratio lower than −2 were called as homozygous deletion.

### Dependency map (DepMap) data analysis

CRISPR dependency data (Dempster *et al*., 2019; Meyers *et al*., 2017)(CERES scores) and gene-level copy number data (Meyers *et al*., 2017) were downloaded from the Public Achilles 2021Q1 DepMap release using the Broad Institute’s DepMap portal. Cell lines were characterized as being ‘*DDX1-MYCN*-coamplified’ if they had DDX1 and *MCYN* copy number value that both were greater than or equal than 2, or ‘*MYCN*-amplified alone’ if they had *MCYN* copy number value that both were greater than or equal to 2 but DDX1 copy number value that was less than 2; cell lines with no copy number data for *DDX1* and *MYCN* were removed from the analysis. From a total cell line in the dependency dataset, 12 were classified as *DDX1-MYCN* co-amplified, and 8 were classified as *MYCN*-amplified. The Wilcoxon rank-sum test was used to compare dependency scores for each gene between the 2 groups. In Figure 3B difference in median gene depletion was plotted on the *x*-axis versus the nominal *P* value of the difference on the *y*-axis. Nominal *P* values are provided. Results of the analysis can be found in a tabular format in the source data.

### Statistical analysis

All experiments were performed a minimum of three times with a minimum of three independent measurements. All statistical analysis was performed with R 3.6 or Python 3.7. All data are represented as mean ± standard error. Statistical significance was defined as *, *P* < 0.05; **, *P* < 0.01, ***, *P* < 0.001.

## Data availability

Copy-number data for 556 neuroblastoma patients were downloaded from https://github.com/padpuydt/copynumber_HR_NB/. Public data of 709 neuroblastoma patients’ microarray supporting the findings of this manuscript were downloaded from ArrayExpress under accession E-MTAB-1781. Cancer cell line metabolism dataset was downloaded from DepMap (Li *et al*., 2019). Public drug response dataset (GDSC2) was downloaded from https://www.cancerrxgene.org/. CRISPR dependency data (Dempster *et al*., 2019; Meyers *et al*., 2017)(CERES scores) and gene-level copy number data (Meyers *et al*., 2017) were downloaded from the Public Achilles 2021Q1 DepMap. Pan-Cancer Analysis of Whole Genomes (PCAWG) study (Kim *et al*., 2020) and Tumor Alteration Relevant for Genomics-driven Therapy (TARGET) database (Pugh *et al*., 2013; Van Allen *et al*., 2014) from cbioportal. All other data are available from the corresponding authors upon reasonable request

## Code availability

Code is available at https://github.com/yeebae1118/DDX1_Bei

## Competing interests

A.G.H and R.P.K are co-founders and shareholders of AMZL therapeutics. The other authors have no competing interests to declare.

## Correspondence

Correspondence and requests for materials should be addressed to

## Author Contributions

Y.B. and A.G.H contributed to the study design and collection and interpretation of the data and wrote the manuscript. Y.B., L.B., A.B., M.K., J.P., S.K., R.F-G., D.W., H.D.G, R.X., L.B., M.G., R.R-F., O.A.S, N.W., J.K. and R.C.G. performed the experiments, analyzed data and reviewed this manuscript. L.B., generated transgenic zebrafish. Y.B., and M.K. performed MS-based protein-protein interaction experiment and analysis. R.K., C.S., J.S., A.E., K.H., J.K., A.I.H.H., P.M., and J.D. contributed to study design. A.G.H. led the study design, to which all authors contributed.

## Acknowledgements

We are grateful to Maria Hondele, Markus Landthaler and Nicole Hübener for critical discussions and to Thomas Look (Dana Faber Cancer Institute, Boston, USA) for transgenic zebrafish and Jan-Philipp Junker (Max Delbrück Center, Berlin, Germany) for providing plasmids. A.G.H. is supported by the *Deutsche Forschungsgemeinschaft* (DFG, German Research Foundation) – 398299703. A.G.H. is supported by the *Deutsche Krebshilfe* (German Cancer Aid) Mildred Scheel Professorship program – 70114107. A.I.H.H. is supported by the *KinderLeben Foundation*. L.B. is supported by the *Deutsche Konsortium für Translationale Krebsfoschung (DKTK)*. This project has received funding from the European Research Council (ERC) under the European Union’s Horizon 2020 research and innovation programme (grant agreement No. 949172). This project was supported by Cancer Research UK and the National Institute of Health (398299703, the eDynamic Cancer Grand Challenge). This project was supported by the Berlin Institute of Health (BIH). Computation has been performed on the HPC for Research cluster of the Berlin Institute of Health.

## Notes

### Competing Interest Statement

Anton G. Henssen and Richard Koche are a co-founder of AMZL therapeutics

### Summary of Updates

I changed the competing interest in the manuscript

