## Supplementary Figures for "Amplicon structure creates collateral therapeutic vulnerability in cancer"

**Supplementary files for: Amplicon structure creates collateral therapeutic vulnerability in cancer**

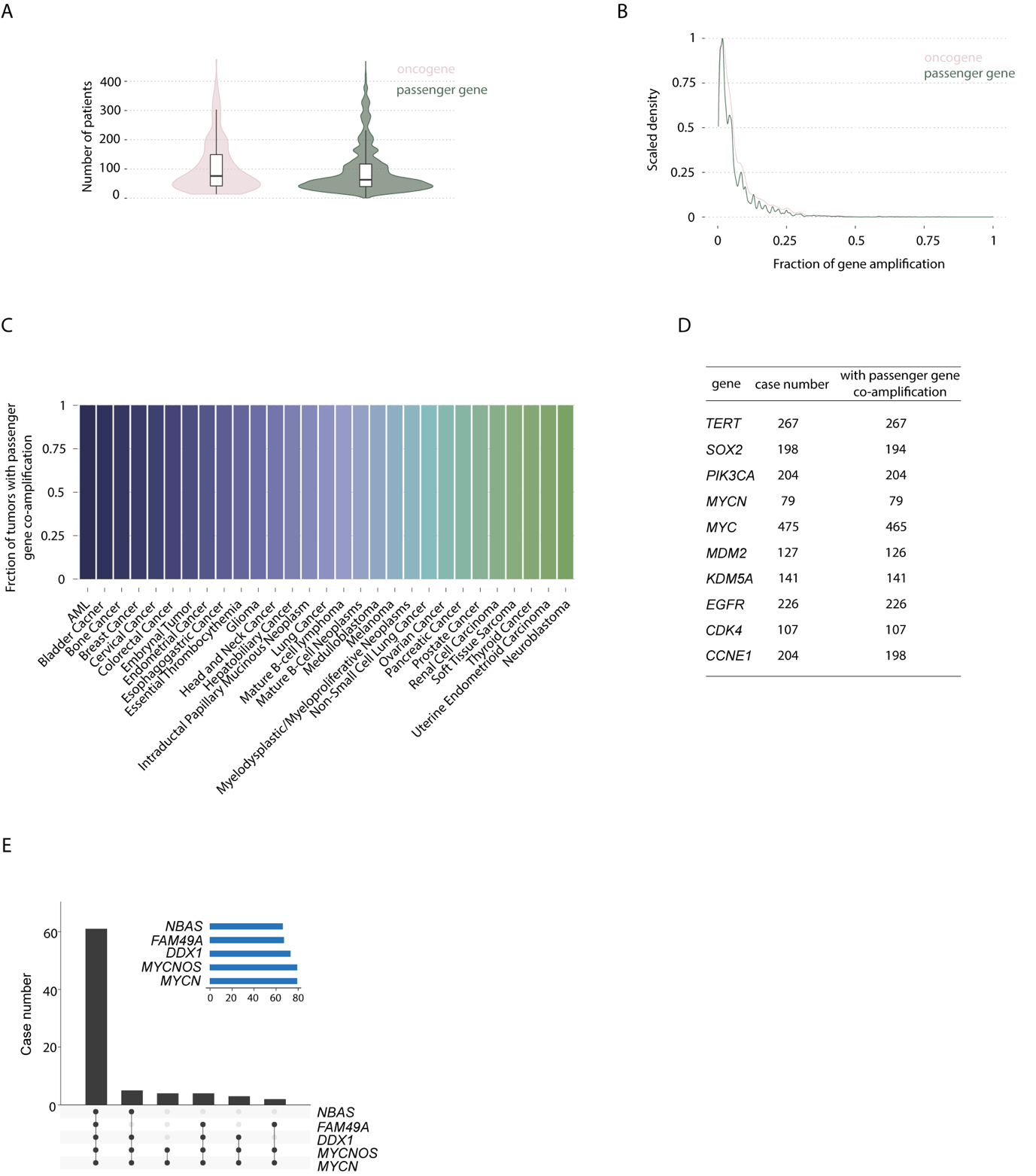

**Supplementary Figure 1. Passenger genes are frequently co-amplified with oncogenes in cancers. A,** Violin plot representing the distribution of patient numbers with oncogene and passenger gene amplifications, respectively (*N* = 2970 total number of cancer patients). **B,** Histogram of the fraction of tumors with oncogene and passenger gene amplification in each tumor entity (*N* = 2970 total number of cancer patients). **C,** Fraction of patients with oncogene amplification also harboring a passenger gene amplification. **D,** 10 representative amplified oncogenes and number of patients with passenger gene co-amplification. **E,** Upset plot showing the co-amplification patterns of 4 passenger genes i.e., *MYCNOS*, *NBAS*, *DDX1* and *FAM49A,* identified in the *MCYN* amplicon and presenting the case number harboring each passenger gene as well as *MYCN* from PCAWG and TARGET dataset.

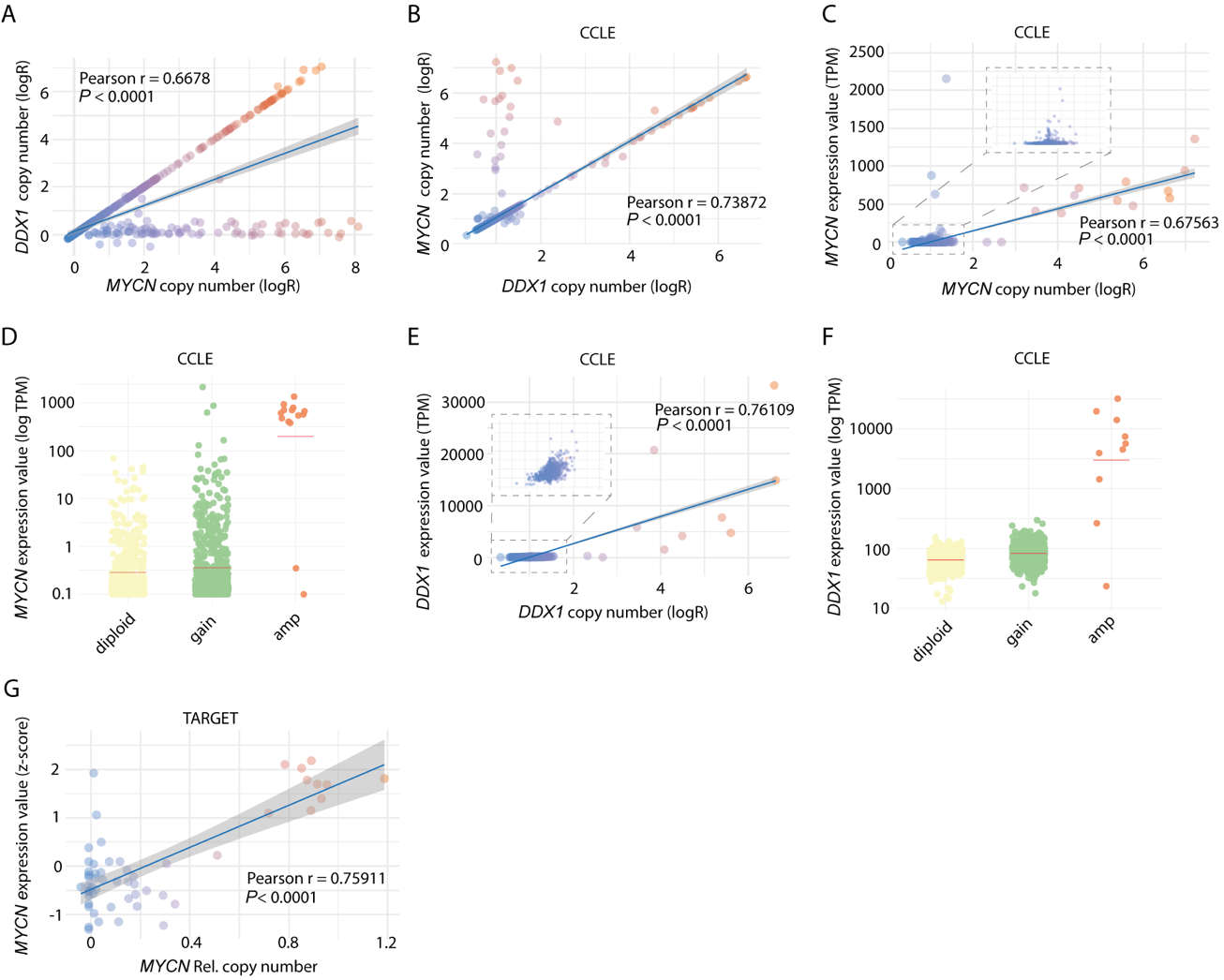

**Supplementary Figure 2. *DDX1* is highly expressed when co-amplified with *MYCN*.**

**A,** *DDX1* copy number compared to *MYCN* copy number in a cohort of 556-neuroblastomas (*N* =556, Pearson r = 0.6678, *P* < 0.0001, *N* = 556). **B,** *DDX1* copy number compared to *MYCN* copy number in cancer cell lines from the CCLE database (*N* = 1713, Pearson r = 0.73872, *P* < 0.0001). **C,** Correlation between *MYCN* copy number and *MYCN* mRNA expression in cancer cell lines from the CCLE database (*N* = 1020, Pearson r = 0.67563, *P* < 0.0001). **D,** *MYCN* mRNA expression in cancer cell lines from the CCLE database (*MYCN* diploid refers to a copy number logR of 1, *MYCN* gain refers to copy numbers logR >1 and <2 and *MYCN* amplification refers to copy numbers of >2). **E,** Correlation between *DDX1* copy number and *DDX1* mRNA expression in cancer cell lines from the CCLE database (*N* = 1020, Pearson r = 0.76109, P < 0.0001). **F,** *DDX1* mRNA expression in in cancer cell lines from the CCLE database, i.e., *DDX1* diploid vs. *DDX1* gain vs. *DDX1* amplification. **G,** *MYCN* copy number compared to MYCN expression in primary neuroblastomas from the TARGET dataset (*N* = 59, Pearson r = 0.75911, P < 0.0001,).

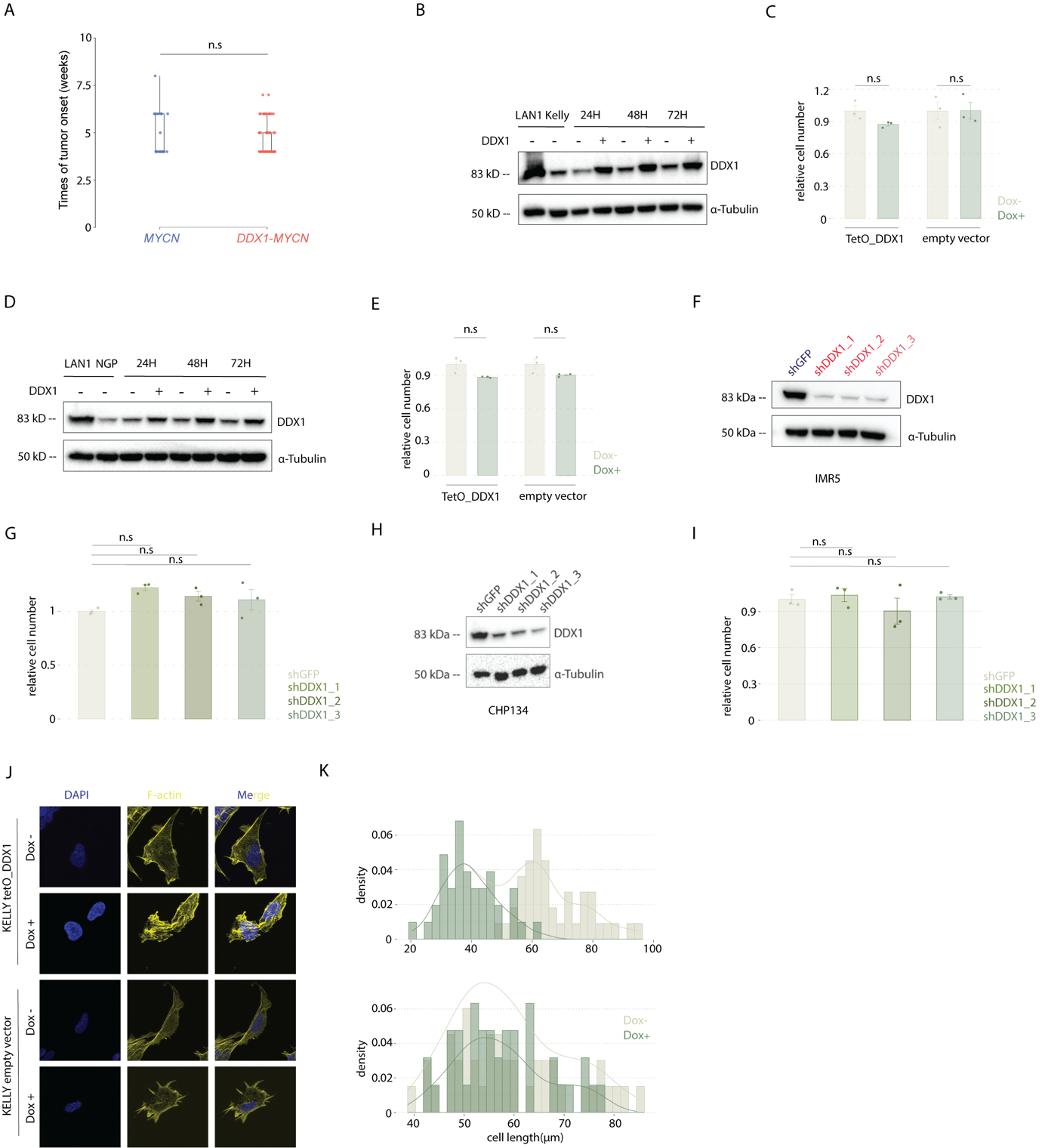

**Supplementary Figure 3. DDX1 expression does not affect tumorigenic properties of cancer cell lines but induces changes in cell size. A,** Time of neuroblastic tumor initiation in MYCN-expressing vs. DDX1-MYCN co-expressing transgenic zebrafish (Welch’s t-test, *P* = 0.783). **B,** Western immunoblot of DDX1 in KELLY cell after inducible expression of DDX1 (1000ng/ml doxycycline treatment for 24, 48 and 72 hours) with LAN1 serving as a positive control and α-tubulin as loading control. **C,** Relative number of viable KELLY cells after inducible expression of DDX1 for 7 days. (Welch’s t-test, *P* = 0.1175 and 0.9852 for cells transduced with an empty vector and cells with an inducible vector treated with doxycycline or vehicle control, respectively). Data are shown as mean ± s.e. (*N* = 3). KELLY cell transduced with an empty vector served as a negative control. **D,** Western immunoblot of DDX1 expression in NGP cell after inducible expression of DDX1 (1000ng/ml doxycycline treatment for 24, 48 and 72 hours) with LAN1 serving as a positive control and α-tubulin as loading control. **E,** Relative number of viable NGP cells after inducible expression of DDX1 for 7 days. (Welch’s t-test, *P* = 0.1061 and 0.1472 for cells transduced with an empty vector and cells with an inducible vector treated with doxycycline or vehicle control, respectively). Data are shown as mean ± s.e. (*N* = 3). NGP cells transduced with an empty vector served as a negative control.

**F,** Western immunoblot of DDX1 in IMR5/75 cells transduced to express shRNAs targeting DDX1 as well as an shRNA targeting GFP serving as a negative control. **G,** Relative number of viable IMR5/75 cells expressing shRNA targeting DDX1 for 7 days compared to cells expressing a shRNA targeting GFP. (Pairwise t-test adjusted by Benjamini-Hochberg correction, *P* = 0.14, 0.31 and 0.31 for all shRNAs, respectively). Data are shown as mean ± s.e. (*N* = 3). **H,** Western immunoblot of DDX1 in CHP134 cells expressing shRNAs targeting DDX1 as well as an shRNA targeting GFP serving as a negative control. **I,** Relative number of CHP134 cells expressing shRNAs targeting DDX1 for 7 days compared to cells expressing a shRNA targeting GFP. (Pairwise t-test adjusted by Benjamini-Hochberg correction, *P* = 0.88, 0.64 and 0.88 for all shRNAs, respectively). Data are shown as mean ± s.e. (*N* = 3). **J,** Representative immunofluorescence images of KELLY cells after inducible expression of DDX1 (1000ng/ml doxycycline treatment for 48 hours). Nucleus and actin cytoskeleton were stained with DAPI (blue) and phalloidin (yellow), respectively. Scale bar: 12 µm. KELLY cells transduced with an empty vector served as negative control. **K**, Histogram of the cellular length of KELLY cells after inducible expression of DDX1 (1 µg/ml doxycycline treatment for 48 hours). KELLY cells transduced with an empty vector served as negative control.

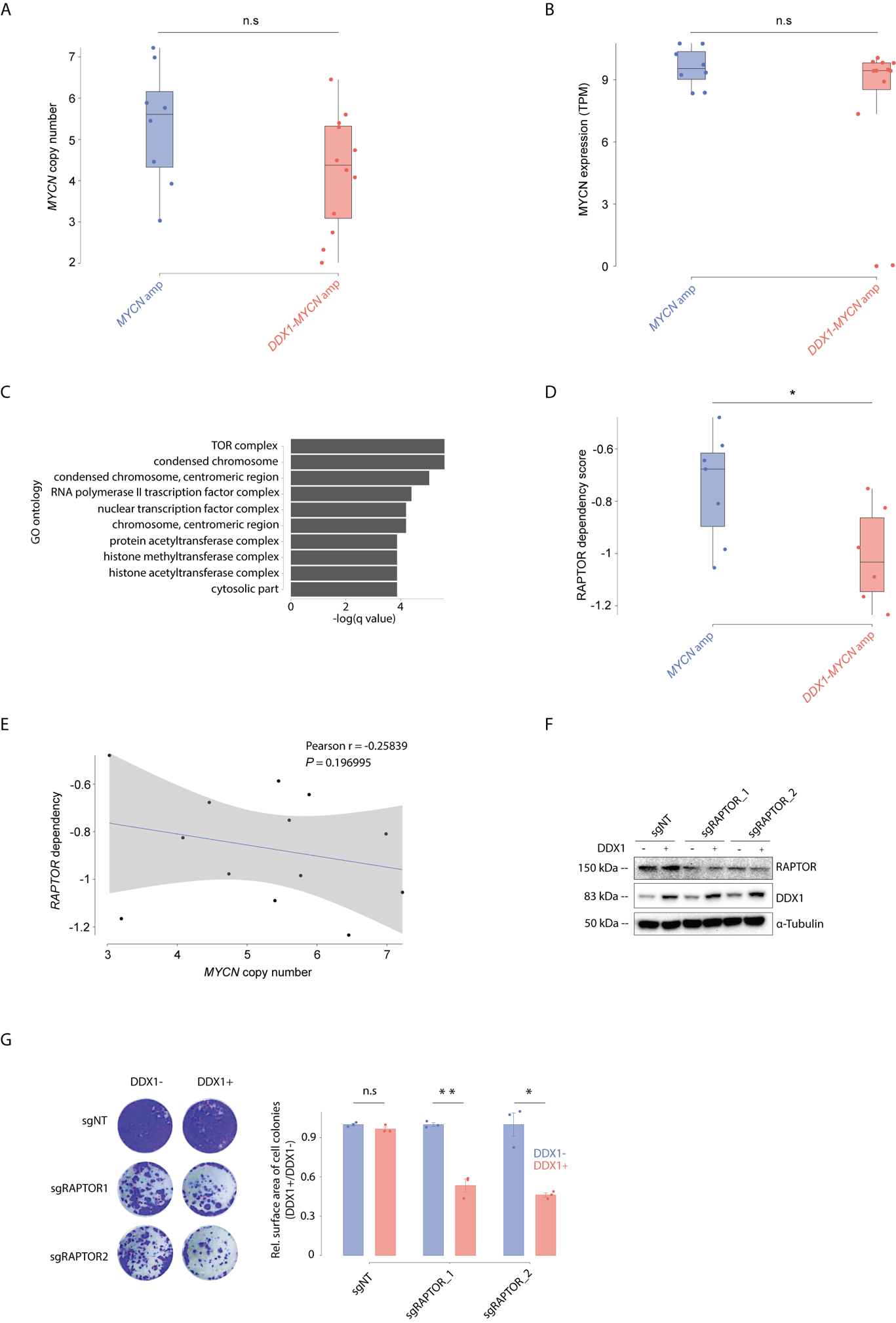

**Supplementary Figure 4. Neuroblastoma cell lines with *DDX1-MYCN* co-amplification depend on mTORC1. A**, Boxplot presenting the distribution of *MYCN* copy number between *MYCN* amplified (*N* = 8) and *DDX1-MYCN* co-amplified (*N* = 12) cancer cell lines**.** Statistical analysis was performed by Wilcox test (*P* = 0.1153). **B,** Boxplot presenting the distribution of MYCN expression between *MYCN* amplified (*N* = 8) and *DDX1-MYCN* co-amplified (*N* = 12) cancer cell lines**.** Statistical analysis was performed by Wilcox test (*P* = 0.4727). **C,** Top 10 GO ontology from GO analysis based on the all-candidate target genes. **D,** Boxplot presenting the distribution of *RAPTOR* dependency score between *MYCN* amplified (*N* = 7) and *DDX1-MYCN* co-amplified (*N* = 6) neuroblastoma cell lines**.** Statistical analysis was performed by Wilcox test (*P* = 0.02564). **E,** Correlation between *MYCN* copy number and the CRISPR-based dependency score from DepMap for *RAPTOR* in neuroblastoma cell lines (Pearson correlation analysis, Pearson r = -0.258396, *P* = 0.19699, *N* = 13). **F,** Western immunoblot analysis of RAPTOR in NGP cells after induced DDX1 expression and incomplete RAPTOR knockout by two independent sgRNAs targeting *RAPTOR* compared to a non-targeting sgRNA. **G,** Representative images of cell colonies formed by NGP cells transduced with the doxycycline-inducible DDX1-mcherry vectors and with two pairs of sgRNA targeting *RAPTOR (*sgRAPTOR*)* or non-target sgRNA (sgNT) as well as Cas9 in the presence and absence of doxycycline (1 µg/ml) and stained with crystal violet (left). **H**. Quantification of colony numbers (right, mean ± s.e. *N* = 3 biological replicates). (Welch’s t-test, *P* = 0.3152, 0.007584 and 0,02257 for sgNT, sgRAPTOR_1 and sgRAPTOR_2, respectively).

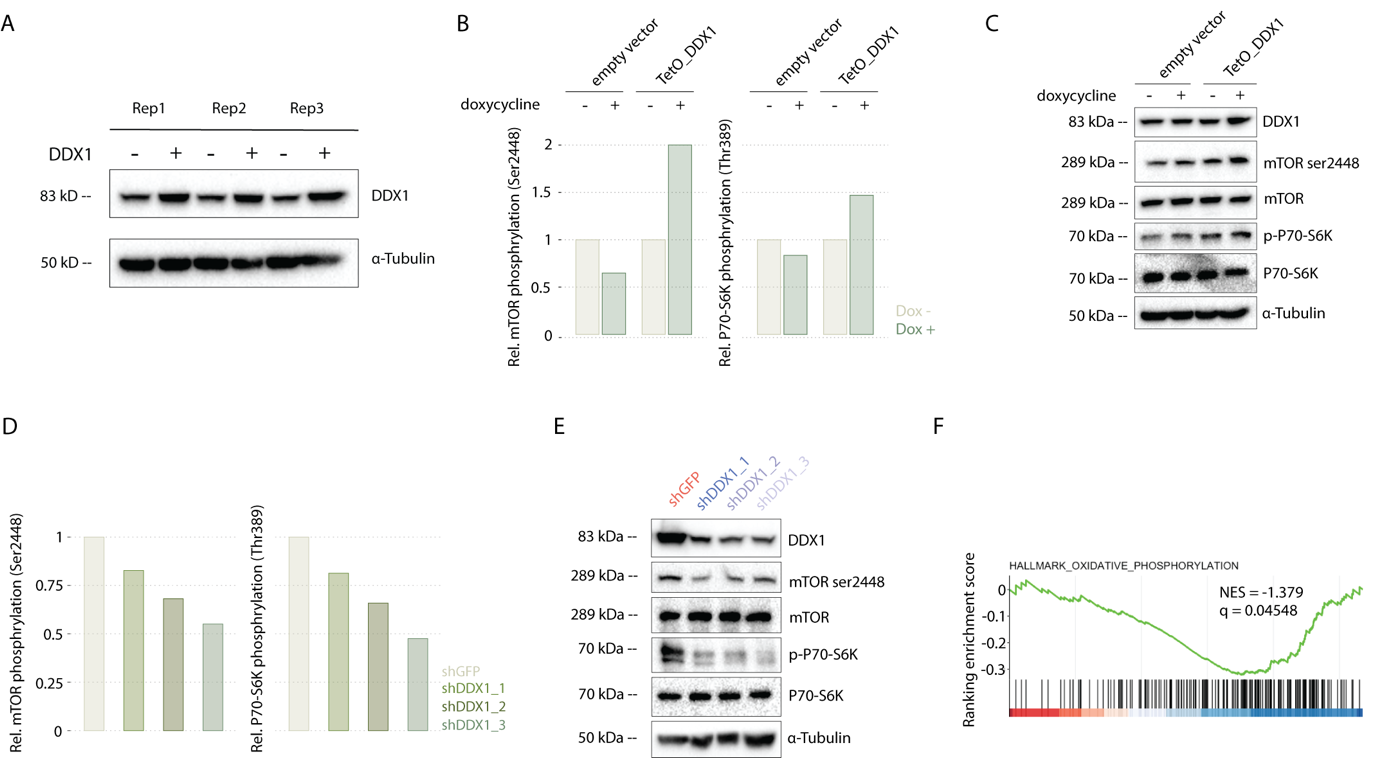

**Supplementary Figure 5. Aberrant DDX1 overexpression results in mTOCR1 pathway activation**. **A,** Western immunoblot of DDX1 in KELLY cells after inducible expression of DDX1 (1000ng/ml doxycycline treatment for 48 hours) with three independent biological replicates. **B,** Quantification of the relative protein expression of mTOR ser2448 phosphorylation and P70-S6K Thr389 phosphorylation in KELLY cell after inducible expression of DDX1 (1000ng/ml doxycycline treatment for 48 hours) as shown in Figure 4E. **C,** Western immunoblot of mTOR ser2448 phosphorylation and P70-S6K Thr389 phosphorylation in NGP cells after inducible expression of DDX1 (1000ng/ml doxycycline treatment for 48 hours). NGP cells transduced with an empty vector served as negative control. **D,** Quantification of mTOR phosphorylation at ser2448 and P70-S6K phosphorylation at Thr389 in IMR5/75 cells expressing shRNAs targeting DDX1 compared to cells expressing a shRNA targeting GFP based on the immunoblot shown in Figure 4F*.* **E,** Western immunoblot mTOR ser2448 phosphorylation and P70-S6K Thr389 phosphorylation in CHP134 cells expressing shRNAs targeting DDX1 compared to cells expressing a shRNA targeting GFP*.* **F,** GSEA based on a set of genes regulating OXPHOS measured in genes differentially expressed in KELLY cells with vs. without ectopic expression of DDX1.

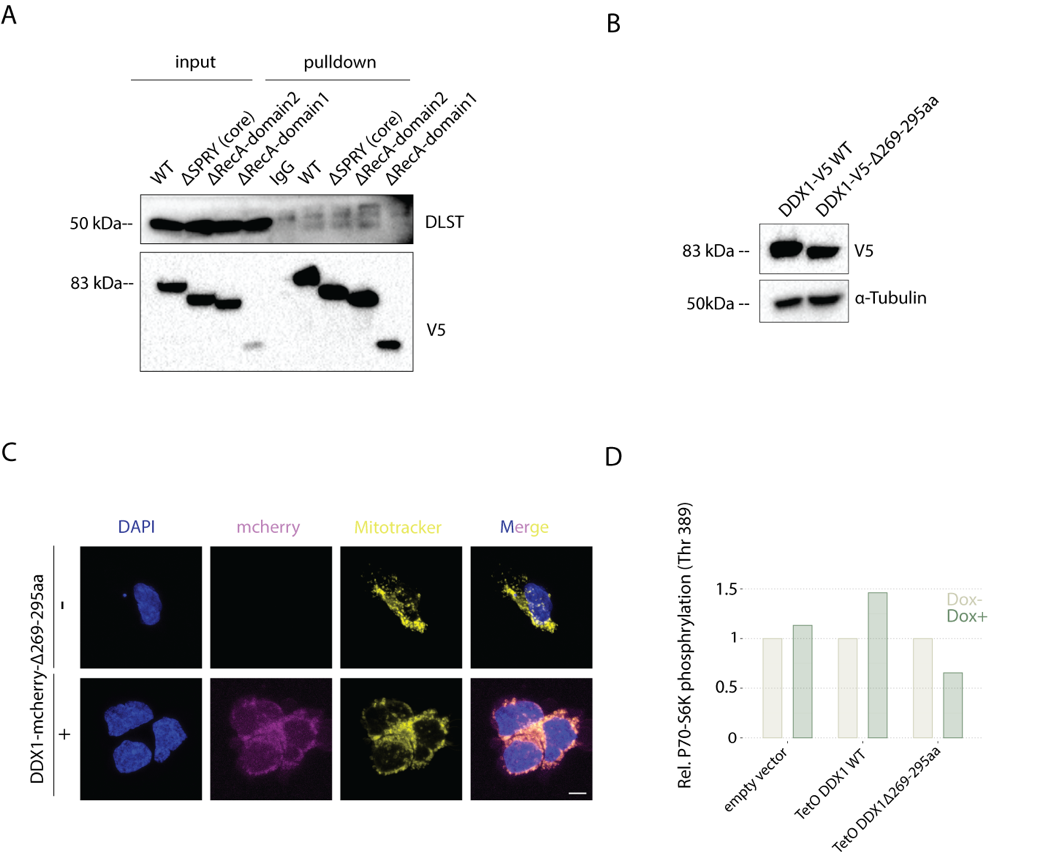

**Supplementary Figure 6. DDX1 interacts with alpha-KGDH complex members and disruption of the DDX1:DLST interaction reduces mTORC1 pathway activation. A,** Western immunoblot of V5, DLST and OGDH before and after immunoprecipitation using antibodies directed against V5, DLST, OGDH or non-specific immunoglobulins (IgG) in KELLY cells expressing DDX1-V5 compared to DDX1-ΔSPRY (core), ΔRecA1 or ΔRecA2 truncation mutants, respectively. **B,** Western immunoblot of DDX1 in NGP cell after inducible expression of DDX1-V5 compared to DDX1-V5-Δ269-295aa (1000ng/ml doxycycline treatment for 48 hours). NGP cell transduced with empty vector serve as controls. **C,** Representative confocal fluorescence imaging photomicrographs of KELLY cells inducibly expressing DDX1-mCherry-Δ269-295aa (magenta), in which mitochondria were stained by MitoTracker DeepRed (yellow) and the nucleus is stained by DAPI (blue; scale bar: 6µm). **D,** Quantification of P70-S6K Thr389 phosphorylation after inducible expression of DDX1 vs. DDX1-V5-Δ269-295aa in KELLY cells (1 µg/ml doxycycline treatment for 48 hours) as shown in Fig. 5K. KELLY cell transduced with empty vector serve as a control.

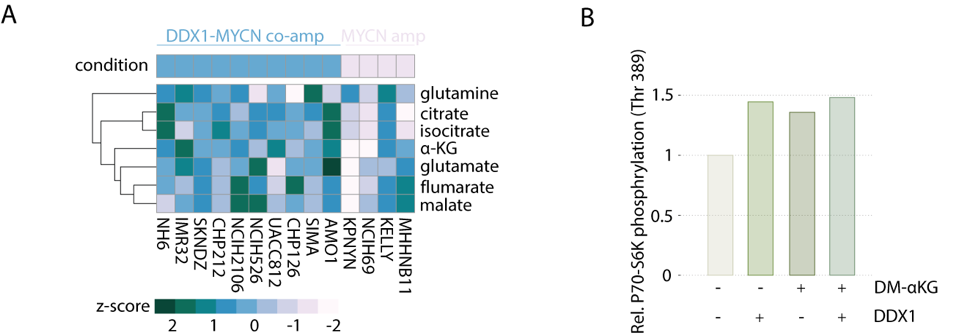

**Supplementary Figure 7. High DDX1 expression is associated with** α**-KG accumulation and OXPHOS reduction. A,** Heatmap of the relative glutamine, citrate, isocitrate, α-KG, glutamate, fumarate and malate concentrations in cancer cell lines with *DDX1*-*MYCN* co-amplification vs. cells with *MYCN* amplifications alone. Cancer cell line metabolism dataset was downloaded from DepMap. **B,** Quantification of P70-S6K Thr389 phosphorylation after incubation of KELLY cells with DM-αKG (2 mM for 48 hours) and inducible DDX1 expression.

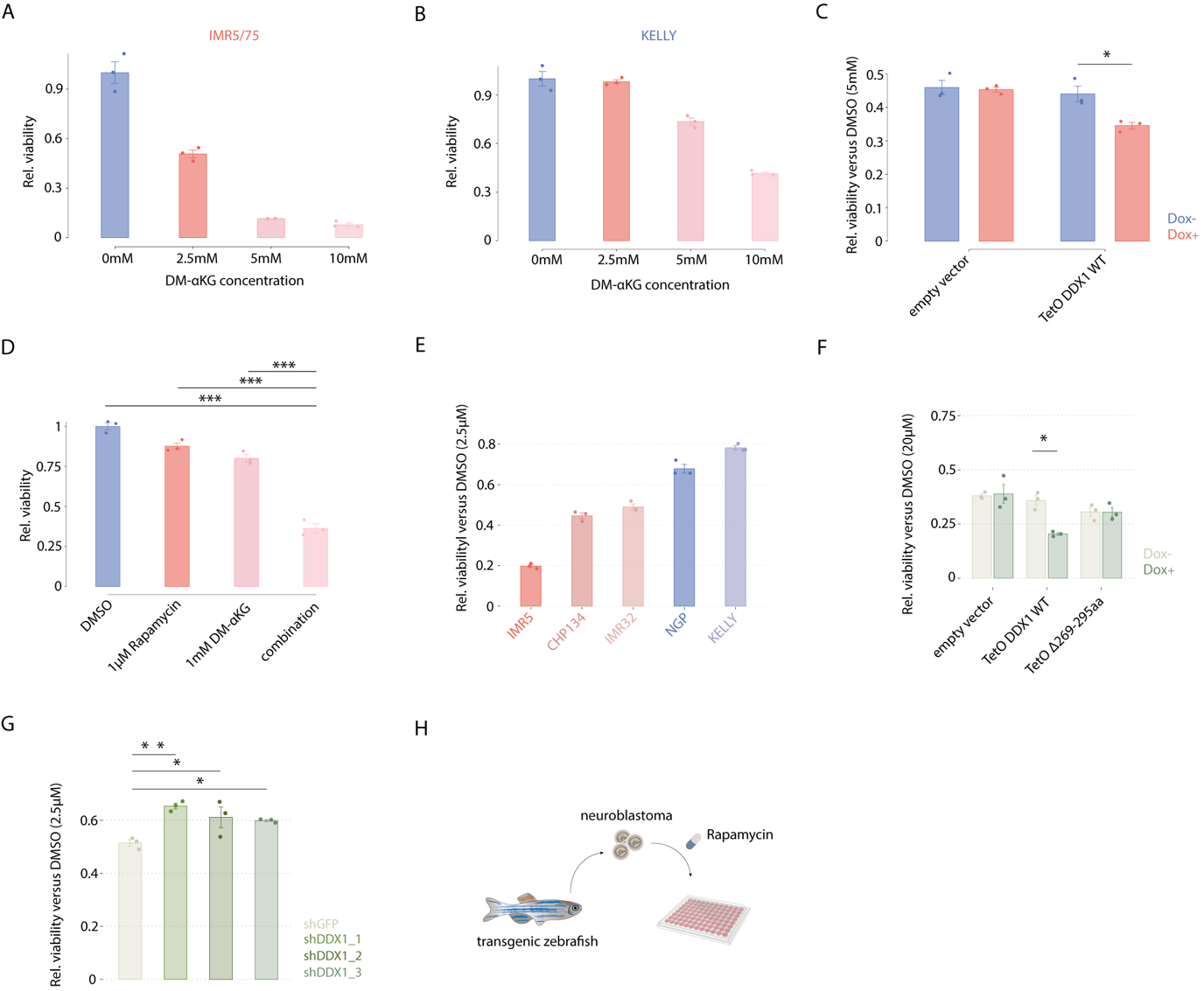

**Supplementary Figure 8. Aberrant DDX1** **expression is associated with increased sensitivity to αKG and pharmacological mTORC1 inhibition. A**, Relative cell viability as measured using MTT assay of IMR5/75 cells harboring a *DDX1-MYCN* co-amplification treated with different concentrations of DM-αKG (0, 2.5, 5 and 10 mM, 72 hours). **B**, Relative cell viability as measured using MTT assay of KELLY cells only harboring a *MYCN* amplification treated with different concentrations of DM-αKG (0, 2.5, 5 and 10 mM, 72 hours)

**C**, Relative cell viability as measured using MTT assay of KELLY cells expressing DDX1 compared to KELLY cells transduced with an empty vector after treatment with DM-αKG (5mM for 72 hours; Welch’s t-test, *P* = 0.03972; Data are shown as mean ± s.e., *N* = 3). **D,** Relative cell viability as measured using MTT assay of IMR5/75 cells after treatment with DM-αKG (1mM) alone, rapamycin (1µM) alone or combination of both compared to DMSO vehicle control treated cells (Welch’s t-test; *P* =2.0e-7, 8.8e-7 and 3.3e-7 for DM-αKG, rapamycin or combination, respectively). Data are shown as mean ± s.e. (*N* = 3). **E**, Relative cell viability as measured using MTT assay of neuroblastoma cell lines with *DDX1*-*MYCN* co-amplification compared to cells with *MYCN* amplification alone after treatment with rapamycin (2.5µM for 72 hours). **F**, Relative cell viability as measured using MTT assay of NGP cells expressing DDX1 compared to NGP cells expressing DDX1 Δ269-295aa after treatment with rapamycin (2.5µM for 72 hours). NGP cells transduced with an empty vector served as negative control. (Welch’s t-test, *P* = 0.01592; Data are shown as mean ± s.e., *N* = 3). **G**, Relative cell viability as measured using MTT assay of CHP134 cells expressing shRNAs targeting DDX1 or GFP (negative control) after treatment with rapamycin (2.5µM for 72 hours). (Pairwise t-test adjusted by Benjamini-Hochberg correction, *P* = 0.01, 0.037 and 0.046 for the three independent shRNAs, respectively; Data are shown as mean ± s.e., *N* = 3). **H**, Schematic of the rapamycin treatment in allografts of transgenic neuroblastic tumors derived from zebrafish.
